# Insular Traveling Waves Link Distributed Neural Dynamics to Human Memory Performance

**DOI:** 10.1101/2025.09.15.676376

**Authors:** Anup Das, Chengyuan Wu, Sameer A Sheth, Joshua Jacobs

## Abstract

The insula is a critical brain region that plays a foundational role in adaptive human behaviors, with diverse subregions performing distinct functional roles. However, explaining how these insular subregions interact to support behaviors is elusive. Using direct recordings from humans performing a spatial episodic memory task, we show that traveling waves within the insula modulate neuronal interactions across insula subregions, by propagating in distinct spatial patterns during specific phases of memory. In addition to traveling plane waves, insula waves also propagated in complex, heterogenous spatial patterns across task conditions. Insular traveling waves correlated with memory success, highlighting the critical role of insular traveling waves in orchestrating memory performance. Our study suggests that insular traveling waves are a key mechanism for modulating interactions and neural coding across regions to support memory processing and potentially a biomarker for investigating dysfunctions in neurological disorders.

## Introduction

The insula is a remarkably important and diverse brain structure, as it underlies cognitive, emotional, and affective control, as well as adaptive human behaviors and consciousness (Menon & Uddin, 2010). Insula dysfunction underlies many psychiatric and neurological disorders (Sha et al., 2019). Over the last three decades, human functional magnetic resonance imaging (fMRI) studies have demonstrated insula involvement in detection and attentional capture of goal-relevant stimuli and flexibly communicating with other large-scale brain networks across a wide range of cognitive tasks (Cai et al., 2016; Dosenbach et al., 2008; Sridharan et al., 2008). Despite its critical role in cognitive control, the specific functional role of the insula in human memory processing, and its electrophysiological foundations, remain elusive.

The subregions of the insula have distinct functions (**Figure 1B**), morphologically, the insula consists of the anterior, middle, and posterior short gyri, as well as separate long gyri around the central sulcus (**Figure 1D**). Cytoarchitectonic analyses of the primate insula have shown that the anterior insula and posterior insula also have different granular structures (Menon, 2025).

**Figure 1:**
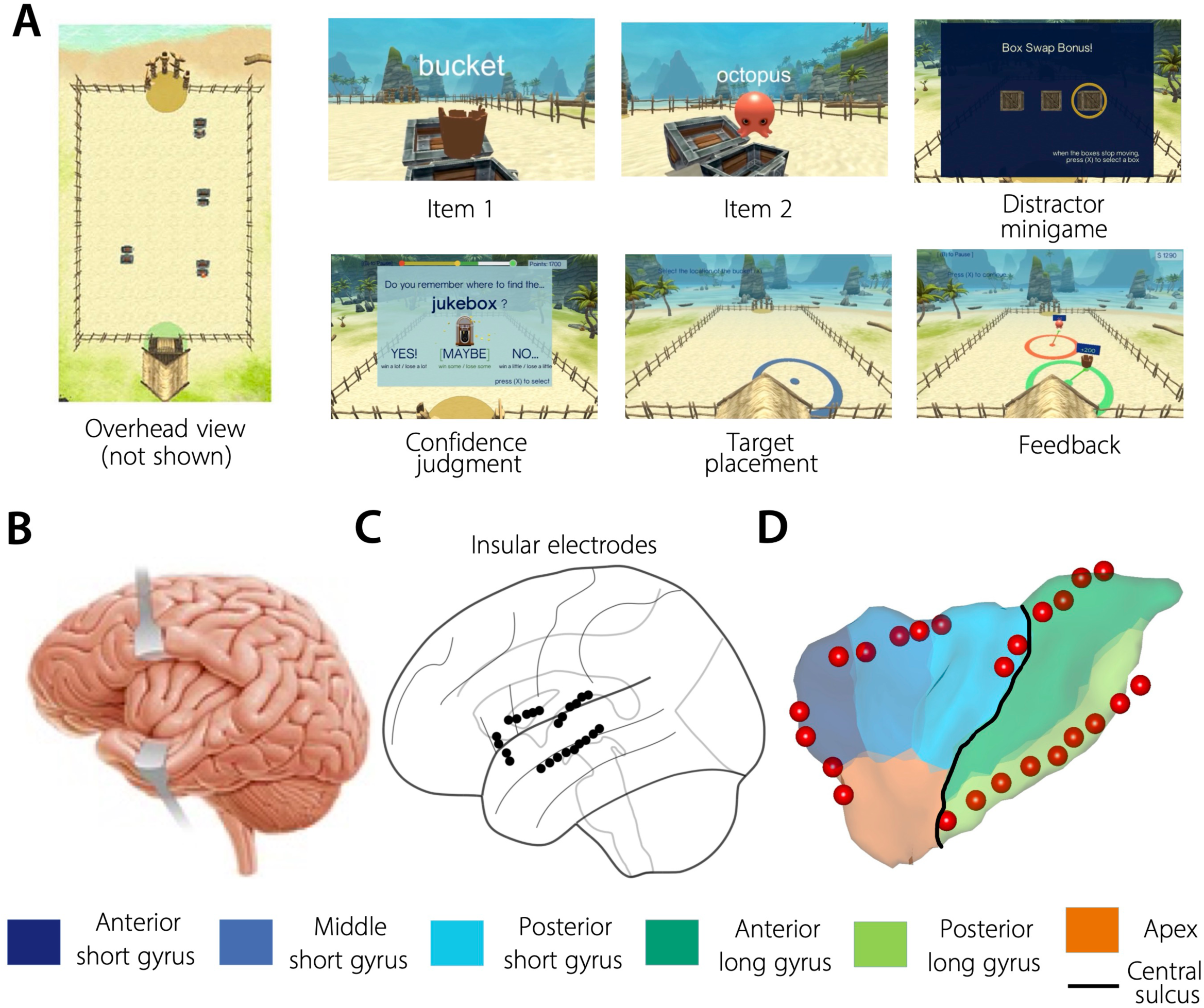
Task structure and insular electrodes. **(A) Spatial episodic memory task.** Patients #1-10 (**Supplementary Figure 1**) performed multiple trials of a Treasure Hunt spatial memory task where they navigated to treasure chests located in a virtual environment containing various objects and after a short delay, were asked to retrieve the spatial location of the objects (**Methods**). **(B) Insula inside the folds of the cerebral cortex. (C) Insular depth electrodes in patient #6, projected on a brain surface plot. (D) Magnifying version of insular electrodes shown in C, overlayed on the different subgyri of the insula.**

Corroborating this anatomical parcellation, fMRI studies have also shown different task-related activations across the insula’s anterior-posterior axis, with the anterior insula supporting high-level cognitive processing and executive functions, whereas the posterior insula integrating sensory information from other cortical areas and forwarding them to the anterior insula (Menon & Uddin, 2010). High level cognition necessarily requires the coordination between the task and sensory processing supported by the anterior and posterior insula, however, the mechanisms governing the interactions between the insula’s functionally and anatomically distinct subdivisions remain unknown.

We propose that the interactions between different subregions of the insula are guided by “traveling waves”, neural activity that progressively propagates across the cortex in specific directions. Traveling waves in other cortical regions correlate with cognitive tasks (Zhang et al., 2018), and we likewise propose that they coordinate activity across insular subdivisions to support high-level cognition and memory. Traveling waves are a key mechanism that can support neuronal computation by selectively and dynamically linking distributed cortical areas, so that certain spatial arrangements of neurons are active at similar times. This phenomenon may be relevant for the insula by allowing the given insular regions that are relevant for a given task to interact (Das et al., 2025). Because propagating oscillations correlate with underlying neural activity (Jacobs et al., 2007), the spatiotemporal organization of traveling waves at each moment may indicate which insular areas are active and how they correspond to particular cortical computational cognitive processes. We therefore hypothesized that propagation of traveling waves may therefore show how the diverse patterns of connectivity within the insula flexibly adapt to task demands and behaviors.

To test this hypothesis, we used an analytical framework for measuring the spatiotemporal organization of propagating brain oscillations including diverse spatial patterns of traveling waves (Das et al., 2025). This framework not only measures plane waves but also spirals and concentric waves. We then applied this framework to examine insular depth recordings from neurosurgical patients performing a spatial episodic memory task. We show that in addition to plane waves, complex spatial patterns of traveling waves in the insula correlated with particular behaviors in episodic memory processing and, crucially, with memory success. We found novel patterns of spatially diverse traveling waves within the insula that correlate with task performance, showing that propagating waves exhibit complex spatial arrangements that reveal when and how different subregions of the insula interact to orchestrate high-level cognition and successful memory encoding.

## Results

### Spatiotemporally stable insular traveling waves

To probe the role of insular traveling waves in memory formation, we examined depth intracranial EEG (iEEG) recordings in the insula from surgical epilepsy patients as they performed a spatial episodic memory task (**Methods, Figure 1**, **Supplementary Figure 1**). We tested whether insular traveling waves reorganize into different directional patterns to distinguish separate behaviors. To examine this hypothesis, we adopted our previously developed analytical framework for measuring traveling waves in iEEG recordings (Das et al., 2022; Das et al., 2025), quantifying their instantaneous spatial structure, and identifying spatial patterns that differentiate individual behavior. One particularly challenging aspect of measuring traveling waves in this region is that the iEEG signals come from multiple implanted depth probes in the insula (for example, **Figure 1C** shows four depth probes spanning the insula). Our analytical framework overcomes this challenge by combining iEEG signals across depth probes and using a spatially localized clustering algorithm.

We examined the role of insular traveling waves in three frequency bands: theta (2-10 Hz), low beta (12-20 Hz), and high beta (20-28). Recent electrophysiology studies in humans and monkeys suggested these bands underlie memory processing, with potentially differential roles for low versus high beta frequency bands, with the former critical for memory and reaction-time, while the later facilitating increased focal attention (Muller et al., 2016; Nougaret et al., 2024; Spitzer & Haegens, 2017). To test for insula traveling waves in each of these bands, we filtered the iEEG signals of the insular electrodes in these three frequency bands and extracted the instantaneous phases at each electrode and timepoint (**Figure 2A**). To measure the spatial phase pattern across these electrodes, we used a circular–linear regression to identify the areas that exhibit a linear relationship between phase and electrode locations *locally* (Das et al., 2022; Das et al., 2025) (**Methods**). This regression approach also automatically estimates the properties of each wave’s propagation, including its phase velocity and direction at each timepoint.

**Figure 2:**
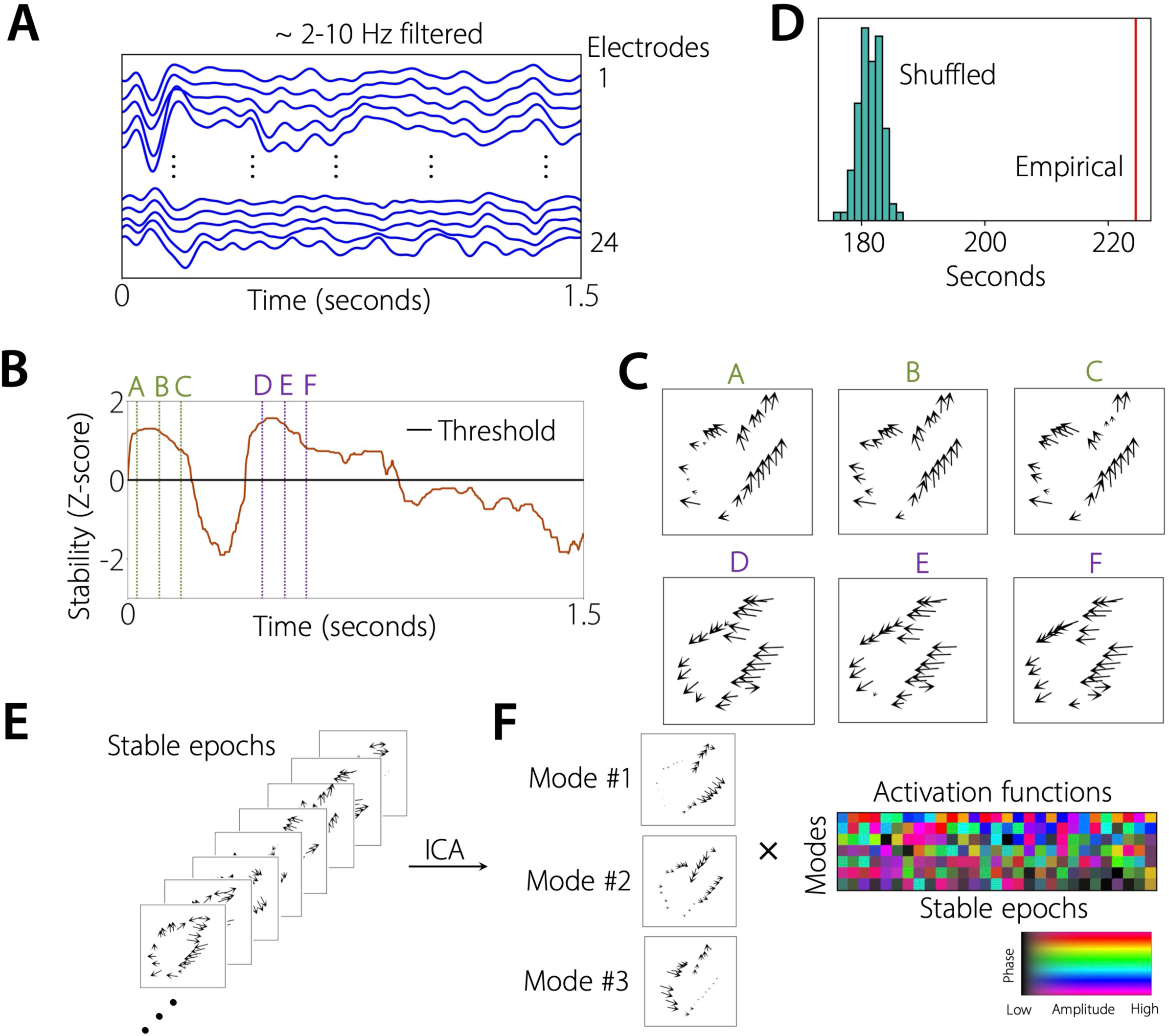
Identifying spatiotemporally stable insular traveling waves. **(A) Filtered signals.** We filtered the signals of the insular electrodes in the theta (2-10 Hz), low beta (12-20 Hz), and high beta (20-28) frequency bands. Shown here are the filtered signals in the 2-10 Hz frequency band from an example trial for the 24 electrodes shown in Figure 1C. Focusing on the peaks (or, troughs) of the filtered signals across electrodes clearly showed the presence of traveling waves. **(B, C) Identifying spatiotemporally stable insular traveling waves.** We used a localized circular-linear regression approach to estimate traveling waves for each time-point and each trial, and in each patient individually (Das et al., 2025) (**Methods**). We then identified stable periods of wave propagation (**Methods**). Shown in **B** are the stability values for an example trial from patient #6. Black line in **B** denotes the stability threshold. In the example trial shown here, there were two stable epochs. Top row in **C** corresponds to the first stable epoch and the bottom row in **C** corresponds to the second stable epoch. Dotted colored vertical green lines in **B** (green in the first stable epoch and purple in the second stable epoch) correspond to time-points for which example traveling waves are shown in **C**. The traveling waves operated in the stable regime for a few tens of milliseconds (top row in **C**), then they entered into the unstable regime where the stable wave pattern broke down and a new wave pattern emerged, and then finally moving onto a new stable regime (bottom row in **C**). Arrows in **C** denote the propagation directions of the waves and lengths of the arrows denote wave strengths. **(D) Shuffling procedure.** We additionally used shuffling procedures as control which suggested that the observed stable epochs are not due to chance (**Methods**). **(E) Independent component analysis (ICA) of traveling waves.** We concatenated the spatial wave patterns across all stable epochs and then passed them as input to ICA (**Methods**). Shown are example stable epochs from patient #6 (2-10 Hz traveling wave). **(F) We extracted the independent activation functions (or, weights) and the corresponding modes as the output from the ICA (Methods).** Activation functions are complex and shown in color with colorbar. Please note that in our previous work, we have extensively validated the application of the ICA procedure to analyzing traveling waves to confirm its sensitivity to extract expected wave patterns and demonstrate specificity to reject negative results in the absence of such patterns (Das et al., 2025).

Recent evidence has shown that traveling waves in humans are very functionally relevant but nonetheless transient (Freeman & Rogers, 2002; Roberts et al., 2019; Schmidt et al., 2023; van Vugt et al., 2007). We thus identified stable periods of insular wave propagation at the single-trial level (**Methods, Figure 2B**) and identified individual epochs with consistent spatial patterns of wave propagation (**Figures 2B, C**). To test the robustness of these stable patterns, we used a shuffling procedure (**Methods**). This analysis revealed that there were reliable spatial patterns within insula traveling waves and that these patterns were statistically robust (all *p’s* < 0.001, **Figure 2D**), ruling out the possibility that they could arise due to chance or noise. The patterns of stability were shorter for spatial patterns in the beta compared to theta frequency band (median 105 msec for low/high beta versus 125 msec for theta, *p* < 0.05, Mann-Whitney U-test).

Consistent with earlier work (Das et al., 2022; Halgren et al., 2019; Zhang et al., 2018), the propagation speed of insular traveling waves was faster for spatial wave patterns at beta versus theta frequencies (median 0.48 m/s and 0.72 m/s for 12-20 Hz and 20-28 Hz respectively versus 0.18 m/s at 2-10 Hz, both *p’s* < 0.001; Mann-Whitney U-tests).

### Extracting the most dominant insular traveling wave patterns using independent component analysis (ICA)

Visually inspecting our results, we observed a diverse range of complex, spatially heterogeneous insular waves in individual subjects at the single trial level. For example, **Figure 2C** shows a spiral traveling wave pattern from a stable epoch in patient #6. The same patient also showed other more complex spatial patterns of traveling waves across other trials (**Figure 2E**).

Moreover, we also saw that, over time, the traveling waves at individual electrodes shifted between different patterns (**Figure 2C**). Interestingly, there were often specific spatial patterns of traveling waves that were distinctive for a given individual. For example, even though spirals and complex waves were the dominant waves in patient #6, patient #10 more often exhibited planar waves and concentric (source/sink) waves (**Figures 4D, 6C**).

**Figure 3:**
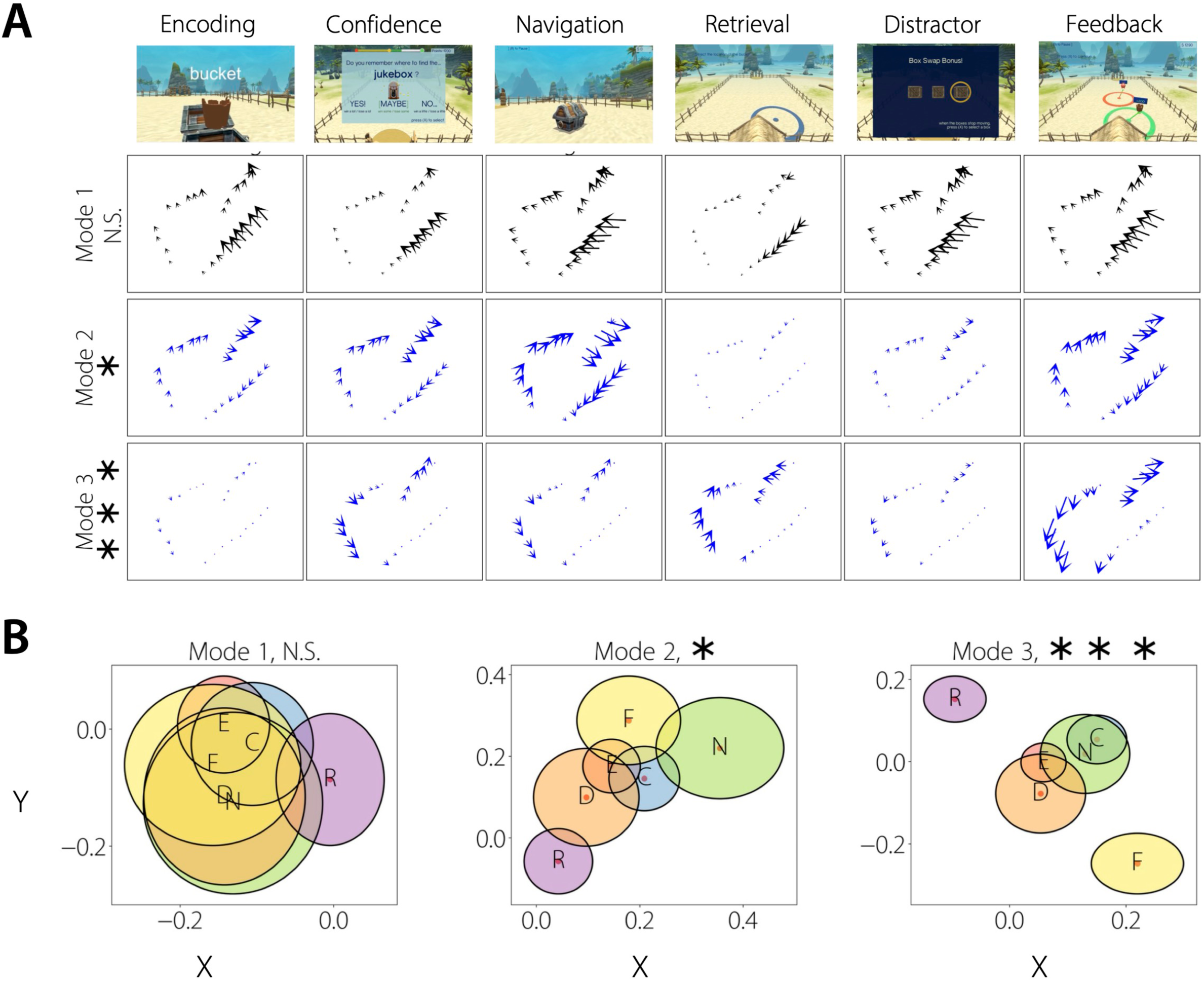
Insular traveling waves can distinguish behavioral states in human memory processing. **(A) Top 3 mean modes of patient #6 (12-20 Hz traveling wave) for each behavioral state.** Mean mode for each behavioral state was calculated as the product of the mean of the activation functions of the mode for the behavioral state with the corresponding mode. Insular traveling waves changed either their direction (for example, retrieval vs. feedback in mode 3), strength (for example, confidence vs. navigation in mode 1), or both (for example, confidence vs. feedback in mode 3), to distinguish behavioral states in the spatial memory task. Statistically significant traveling waves in MANOVA are plotted in blue and non-significant traveling waves are plotted in black. **(B) Distinguishing behavioral states in this patient in the spatial episodic memory task, shown are the activation functions in the complex plane for the three modes in A.** The shift in direction and/or strength of the insular traveling waves between different behaviors can be visualized in terms of the activation functions where, a change in the direction of the waves corresponds to a change in the angle of the activation functions (for example, compare retrieval and feedback for mode 3 in **A and B**), a change in the strength of the waves corresponds to a change in the magnitude/length of the activation functions (for example, compare confidence and navigation for mode 2 in **A and B**), a change in both the direction and strength of the waves corresponds to a change in both the angle and length of the activation functions (for example, compare confidence and feedback for mode 3 in **A and B**). For each ellipse (task condition), the major axis (horizontal axis) denotes the standard-error-of- the-mean (SEM) for the real-part and the minor axis (vertical axis) denotes the SEM for the imaginary part, of the activation functions. E: Encoding (N=363), C: Confidence (N=226), N: Navigation (N=87), R: Retrieval (N=221), D: Distractor (N=100), F: Feedback (N=110). *** *p* < 0.001, * *p* < 0.05 (FDR-corrected).

**Figure 4:**
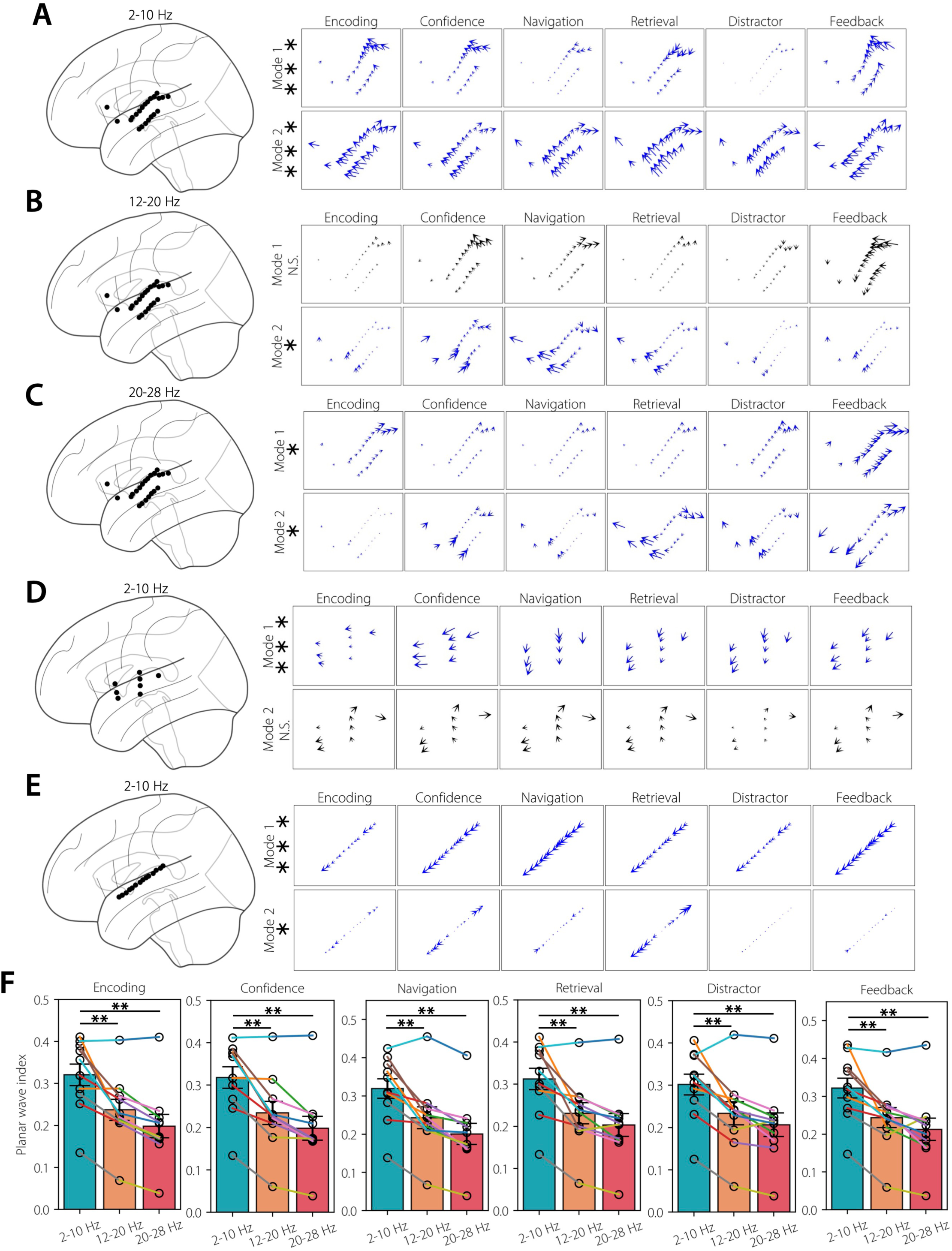
Insular traveling waves can distinguish behavioral states across frequencies and subjects. **(A) 2-10 Hz insular traveling wave in patient #7.** Shown are top 2 mean modes for each behavioral state. N=630 for encoding, N=396 for confidence, N=165 for navigation, N=392 for retrieval, N=156 for distractor, and N=341 for feedback. **(B) Same as in A, but in the 12-20 Hz frequency.** N=540 for encoding, N=326 for confidence, N=130 for navigation, N=342 for retrieval, N=127 for distractor, and N=284 for feedback. **(C) Same as in A, but in the 20-28 Hz frequency.** Note more complex, heterogeneous spatial patterns of traveling waves in low beta and high beta frequencies compared to the theta frequency. N=450 for encoding, N=286 for confidence, N=102 for navigation, N=277 for retrieval, N=115 for distractor, and N=258 for feedback. **(D) Same as in A, but in patient #10.** N=1800 for encoding, N=1120 for confidence, N=459 for navigation, N=1092 for retrieval, N=444 for distractor, and N=817 for feedback. **(E) Same as in A, but in patient #9.** Note that this patient had a single depth probe, and therefore one-dimensional traveling waves. Nevertheless, we found the presence of a source wave in mode 2 of this patient. N=1868 for encoding, N=1140 for confidence, N=461 for navigation, N=1128 for retrieval, N=446 for distractor, and N=1147 for feedback. **(F) Comparing planar wave index across frequencies in each behavioral state.** We obtained lower planar wave indices in the low beta and high beta frequencies compared to the theta frequency, indicating that insular traveling waves take the shape of more complex spatial patterns at higher frequencies compared to lower frequencies. N=10 across behavioral states. *** *p* < 0.001, ** *p* < 0.01, * *p* < 0.05 (FDR-corrected).

**Figure 5:**
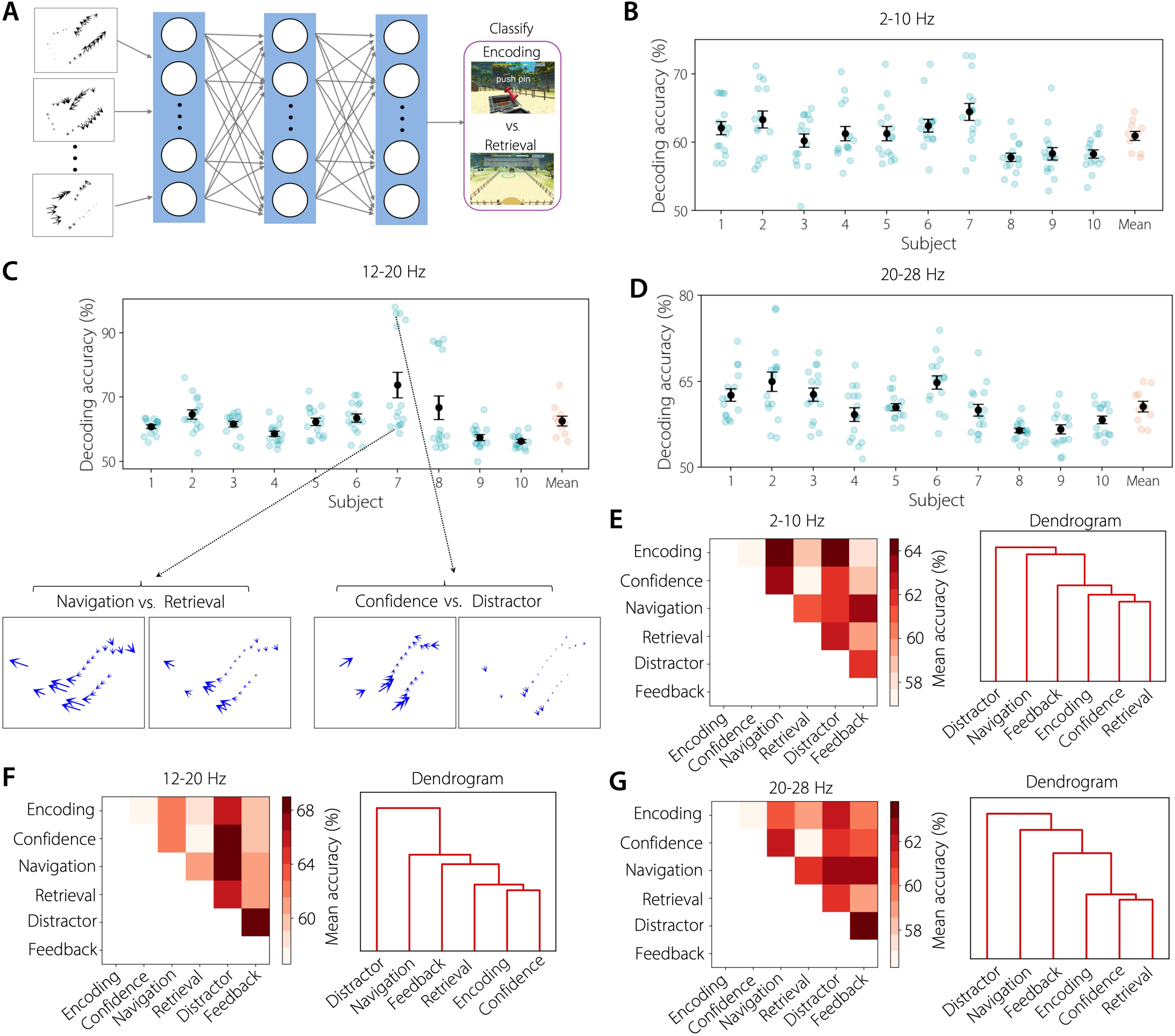
Decoding behavioral states of human memory representations from insular traveling waves. **(A) Neural network architecture.** We used a multilayer neural network with an input layer, three hidden layers (32, 16, and 8 neurons respectively), and an output layer for decoding pairwise behavioral states (for example, encoding vs. retrieval) (see **Methods** for details). We used the extracted weights from the ICA procedure as features for training our neural network classifiers. **(B) Network decoding accuracy in the 2-10 Hz frequency.** Each point in green corresponds to a pair of behavioral states (for example, encoding vs. retrieval) within a subject. Error bars show SEM decoding accuracies across all pairs. Each point in red corresponds to the mean decoding accuracy across all behavioral states for a given subject. Error bars show SEM decoding accuracies across all subjects. Note that chance level is at 50%. **(C) Same as in B, but for the 12-20 Hz frequency.** Pairs of behavioral states with higher decoding accuracy had distinct traveling wave patterns (for example, confidence vs. distractor) compared to those with lower decoding accuracy for which traveling waves were more similar to each other (for example, navigation vs. retrieval). **(D) Same as in B, but for the 20-28 Hz frequency. (E) Mean decoding accuracy across all subjects for each pair of behavioral states.** Dendrogram shows that the navigation and distractor states were the most decodable from the other states, but relatively less decodable from each other. **(F) Same as in E, but for the 12-20 Hz frequency. (G) Same as in E, but for the 20-28 Hz frequency.**

**Figure 6:**
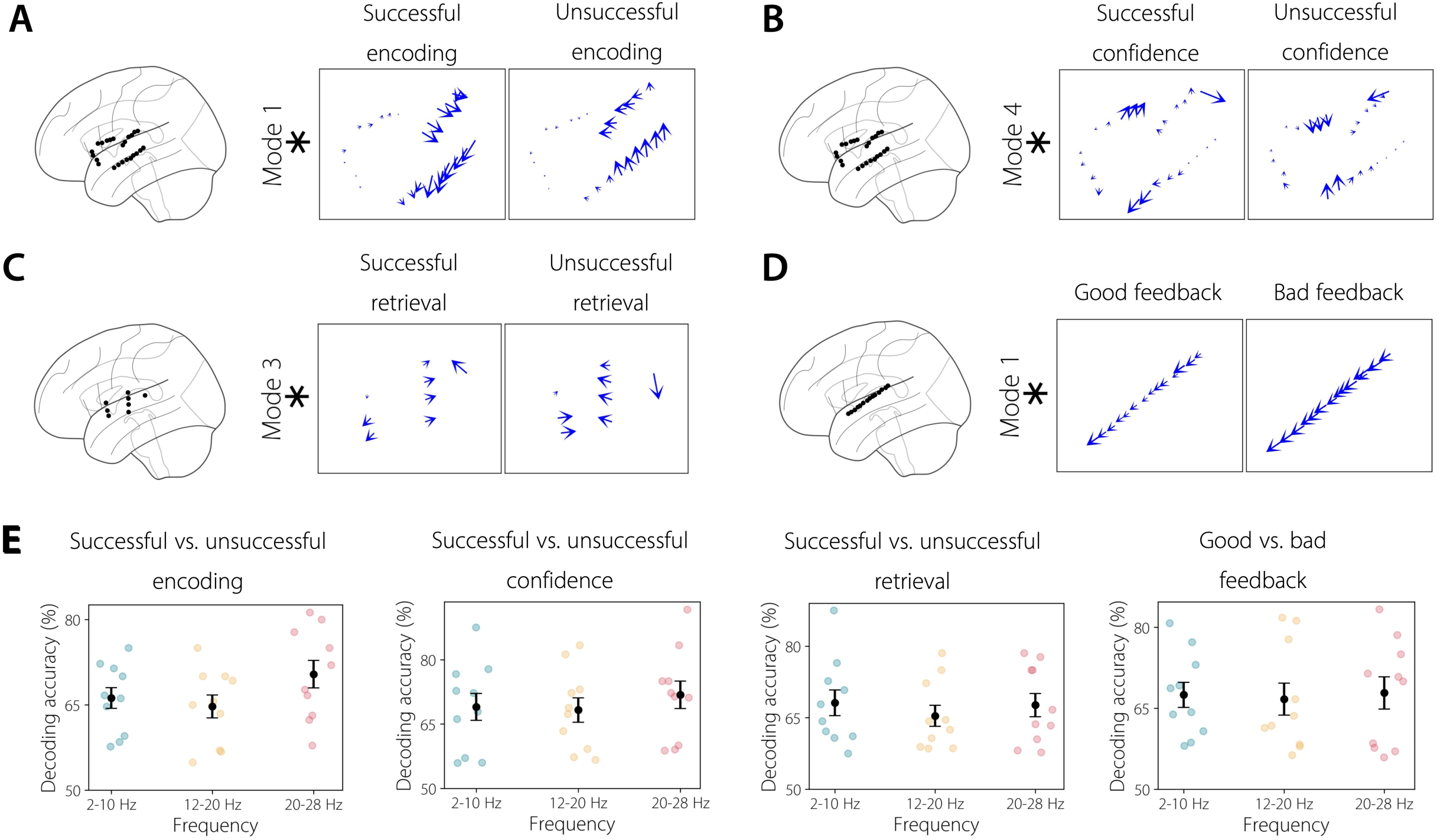
Insular traveling waves correlate with memory performance. **(A)** Insular traveling wave can distinguish successful from unsuccessful memory encoding. Shown are traveling waves in mode 1 in the 2-10 Hz frequency for patient #6. Note the presence of a clockwise spiral during successful memory encoding and the presence of a counter-clockwise spiral when memory encoding was unsuccessful. N=46 for successful encoding, N=219 for unsuccessful encoding. **(B) Insular traveling wave can distinguish successful from unsuccessful confidence.** Shown are traveling waves in mode 4 in the 2-10 Hz frequency for patient #6. Note the presence of a complex spatial pattern of traveling wave during successful memory confidence which changes its shape when memory confidence was unsuccessful. N=43 for successful confidence, N=222 for unsuccessful confidence. **(C) Insular traveling wave can distinguish successful from unsuccessful memory retrieval.** Shown are traveling waves in mode 3 in the 2-10 Hz frequency for patient #10. Note the presence of a source during successful memory retrieval and the presence of a sink when memory retrieval was unsuccessful. N=536 for successful retrieval, N=556 for unsuccessful retrieval. **(D) Insular traveling wave can distinguish good and bad feedback.** Shown are traveling waves in mode 1 in the 2-10 Hz frequency for patient #9. Note the presence of a higher wave-strength traveling wave during good feedback period and the presence of a lower wave-strength traveling wave during bad feedback period. N=710 for good feedback, N=437 for bad feedback. * *p* < 0.05 (FDR- corrected). **(E) Memory performance can be decoded from insular traveling waves.** Panel 1: Decoding accuracies for successful vs. unsuccessful memory encoding in each frequency. Each data point represents one subject. Panel 2: Same as in panel 1, but for successful vs. unsuccessful confidence. Panel 3: Same as in panel 1, but for successful vs. unsuccessful retrieval. Panel 4: Same as in panel 1, but for good vs. bad feedback.

To quantitatively distinguish the full diversity of spatial wave patterns over time in each subject, we used an algorithm based on independent component analysis (ICA) (Fu et al., 2015; Li & Adalı, 2010). This algorithm separately identified the range of spatial patterns of insular traveling waves at each frequency and in each patient (see **Methods**), through the measurement of time-varying activation functions that quantified the waxing and waning of individual spatial traveling wave patterns over time in each patient. Using this algorithm, we labeled each spatial wave pattern, which we refer to as a “mode” (**Figure 2F**, **Methods**) and then measured the magnitude and direction of each pattern for each epoch with the “activation functions”. Thus, by examining the modes and activation functions, we can identify the spatial patterns of traveling waves that were most strongly present at each moment in the recording.

Application of this ICA procedure on our data revealed a wide range of spatial traveling wave patterns across subjects (**Methods, Figures 3, 4**). Individual subjects showed a mean of 3 significant modes, indicating that there were generally multiple spatial patterns of traveling waves in each patient. On average, the first three modes explained ∼53%, ∼21%, and ∼11% variance respectively. Together, these first three modes thus explained ∼ 85% variance in the traveling waves, indicating that reliable spatial patterns explain a large amount of the apparent variance in the arrangements of brain oscillations. Thus, ICA, can identify the diverse insular traveling wave patterns that are present at each frequency, and reveal how the spatial pattern of wave propagation varies throughout the task.

The recent literature emphasized the emergence of complex spatial patterns of traveling waves in the beta band, compared to the predominantly planar waves found at lower frequencies (Das et al., 2025; Muller et al., 2016). Thus, we also examined the complexity of the insular traveling wave patterns at each frequency. Visually, we observed more complex waves in the higher frequency beta bands compared to the lower frequency theta band (**Figures 4A-C**). To quantify this phenomenon, we defined “planar wave index”, which measures the similarity of the direction of wave propagation across electrodes. If the traveling waves for individual electrodes are all propagating in the same direction, i.e., a planar wave, then their vector sum will be a higher value. This analysis revealed that the planar wave indices in both the low beta and high beta frequency bands were significantly lower compared to the theta frequency band across task conditions, indicating the presence of complex spatial patterns of insular traveling waves in higher frequency beta bands (*p’s* < 0.01, Wilcoxon signed-rank tests, **Figure 4F**).

Together, these results suggest an intricate multiplexing of both simpler planar waves and more complex spatial propagation patterns of waves in distinct frequency bands, which putatively enables the insula to flexibly and simultaneously use distinct frequency channels for integrating information across its subdivisions for memory processing.

### Insular traveling waves encode behavioral states of human memory representations

Next, to identify the functional role of insular traveling waves, we examined how the prevalence of traveling waves with different shapes shifted between stages of memory. Insular traveling waves showed different propagation patterns between stages of memory processing (**Figures 3, 4**). As an example, **Figures 3A, B** show an example subject where insular electrodes showed three distinct modes of traveling waves, out of which waves corresponding to modes #2 and #3 changed their propagation direction between the stages of memory (*p’s* < 0.05, MANOVA). We observed a spiral wave pattern in mode #2 which was nearly absent during retrieval and distractor stages (as indicated by its low magnitude) but was strongly present across all other task conditions. Similarly, we observed a more complex spatial wave pattern in mode #3, which was nearly absent during the encoding and distractor periods, exhibited relatively stronger magnitude during the confidence, navigation, and retrieval periods, and strongest during the feedback period.

Our ICA approach can adaptively estimate the different spatial patterns of traveling waves present in different individuals, therefore allowing us to measure the inter-individual differences in traveling waves as well. We indeed observed different spatial patterns of traveling waves across individuals. For example, for subject #7 in **Figure 4A**, we observed that mode #1 was almost absent during navigation and distractor periods, however was strongly present across all other task conditions. For subject #10 in **Figure 4D**, we observed that the planar traveling wave in mode #1 changed its direction across conditions to distinguish behaviors. This is in contrast to the traveling waves in subject #6 in **Figure 3A**, where we observed complex spatial patterns of traveling waves such as spirals. We also observed behavioral state-specific traveling waves where there was a single depth probe in the insula. For example, **Figure 4E** shows a single depth probe in the insula, where in mode #1, we observed a planar traveling wave which was the strongest during navigation, but weakest during encoding. Mode #2 of the same depth probe exhibited a source wave which was the strongest during retrieval, however relatively weaker during other periods of the memory task. These behavioral state-specific insular traveling waves were prominent across all 10 subjects (all *p’s* < 0.05, MANOVA).

Due to the relatively spatially sparse sampling of insular depth probes in individual participants, it was not possible to classify the identified ICA modes into broad spatial propagation patterns of waves such as rotational, concentric (source/sink), etc. (cf. (Das et al., 2025)). Rather, here we focused on the features of these traveling waves and their functional and anatomical relevance.

Therefore, in addition to measuring the spatial patterns of traveling waves in each subject individually, we also examined whether certain features of traveling waves changed across time to distinguish behaviors at the group level (across-subjects). We separately examined how the propagation direction and strength of the traveling waves shifted between task conditions to distinguish behaviors. This analysis revealed that the strength of the traveling waves did not differ between task conditions such as encoding, retrieval, navigation, etc. at the group level in any frequency band (*p’s* > 0.05, Kruskal-Wallis tests). Because of the functional significance of the anterior-posterior parcellation of the insula, we also examined the effect of the anterior-posterior direction of the traveling waves in distinguishing task conditions. However, we did not find any significant effect of the anterior-posterior direction on behavior at the group level in any frequency (*p’s* > 0.05, Kruskal-Wallis tests). Together, these results corroborate our findings above that show that there are specific spatial patterns of traveling waves in a given individual that are functionally relevant for encoding various behaviors and the spatial patterns of these traveling waves vary across subjects, with no specific spatial pattern consistently present across subjects. This also underscores the importance of our ICA approach that can adaptively estimate the spatial patterns of these waves, therefore allowing us to measure the inter-individual differences in traveling waves.

### Insular traveling waves encode memory performance

In addition to examining the functional role of traveling waves in separate memory states, we also examined whether the insular traveling waves also shifted to distinguish performance, i.e., whether the traveling waves shifted their propagation direction and/or strength to distinguish successful compared to unsuccessful memory encoding and retrieval. Insular traveling waves indeed showed different propagation patterns between successful versus unsuccessful memory encoding and retrieval (**Figures 6A, C**). In each recall period, immediately prior to recall, subjects indicated their confidence to remember the location of the object (“high”, “medium”, or “low”), which essentially corresponds to cued retrieval. If subjects were able to retrieve the locations of the objects after indicating high confidence, this constituted successful confidence, whereas failure to retrieve the locations of the objects after indicating high confidence constituted unsuccessful confidence. Insular traveling waves also shifted their propagation patterns between successful versus unsuccessful confidence (**Figure 6B**) as well as good versus bad feedback (**Figure 6D**).

As an example, **Figure 6A** shows a spiral wave in patient #6 in mode #1 where the spiral moved in the clockwise direction during successful memory encoding, but in the counter-clockwise direction when memory encoding was unsuccessful. In the same subject, a complex spatial pattern can be seen in mode #4 during successful confidence, which changed direction during unsuccessful confidence (**Figure 6B**). In another subject, in mode #3, insular traveling waves propagated as a source wave during successful memory retrieval, but propagated as a sink wave when memory retrieval was unsuccessful (**Figure 6C**). Similarly, stronger traveling waves can be observed in an example subject in mode #1 during bad compared to good feedback (**Figure 6D**). These performance specific insular traveling waves were very common as insular electrodes showed significant shifts in traveling wave direction or strength between successful and unsuccessful memory encoding in 9 out of 10 subjects, successful versus unsuccessful memory retrieval in 9 out of 10 subjects, successful versus unsuccessful confidence in 9 out of 10 subjects, and good versus bad feedback in 7 out 10 subjects.

To more broadly understand this phenomenon, we examined at the group level whether the strength and direction of the traveling waves changed across time to correlate with memory performance. This analysis revealed that the strength of the traveling waves increased for successful compared to unsuccessful memory encoding in the low beta band, whereas the opposite was true in the theta band (**Figure 7A**, *p’s* < 0.01, Wilcoxon sign-rank tests). There was a similar frequency dependence during the feedback period where traveling waves increased their strength for good compared to bad feedback in the high beta band, whereas the opposite was true in the theta band (**Figure S2B**, *p’s* < 0.01, Wilcoxon sign-rank tests). Critically, this signature was observed in every single subject, highlighting the robustness of our findings.

**Figure 7:**
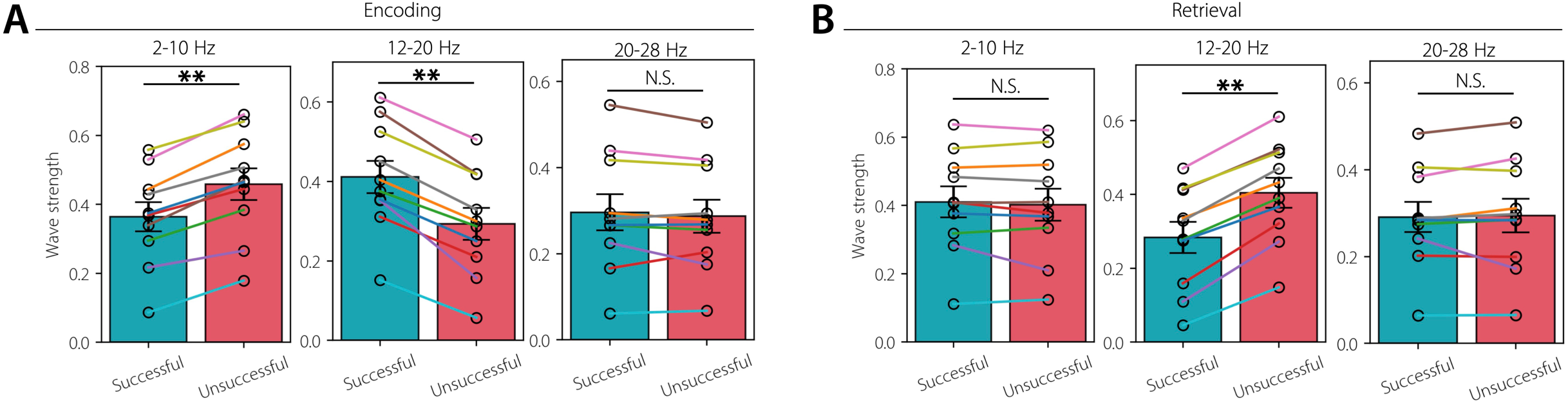
Wave strength of insular traveling waves can distinguish memory performance across subjects. **(A) Wave strength of insular traveling waves for successful vs. unsuccessful memory encoding in each frequency band.** Wave strength increased for successful compared to unsuccessful memory encoding in the 12-20 Hz frequency. This pattern was reversed in the 2- 10 Hz frequency. Error bars denote SEM across subjects. N=10. **(B) Same as in A, but for the retrieval period.** Wave strength increased for unsuccessful compared to successful memory retrieval in the 12-20 Hz frequency. ** *p* < 0.01, N.S. Not significant (FDR-corrected).

Moreover, we found stronger traveling waves during unsuccessful compared to successful memory retrieval in the low beta band (**Figure 7B**, *p* < 0.01, Wilcoxon sign-rank test).

We also considered the possibility that the differential wave strength that we observed for successful compared to unsuccessful memory above, could have been driven by a difference in signal power. Therefore, we carried out control analyses comparing the signal power for successful versus unsuccessful memory encoding, etc. (see **Methods**). This analysis revealed that power does not differ between successful compared to unsuccessful memory encoding, recall, or confidence as well as good compared to bad feedback, in any of the frequencies (*p’s* > 0.05, Wilcoxon sign-rank tests), suggesting that traveling waves, rather than signal power, encodes memory performance.

Similarly, we also examined the effect of the anterior-posterior direction of the traveling waves on memory performance at the group level, however, we did not find any significant effect (*p’s* > 0.05, Wilcoxon sign-rank tests). Together, these results indicate that wave strength is a stronger correlate of memory performance compared to wave direction, in the human insula.

### Memory states can be decoded from insular traveling waves at the single trial level

Since the spatial patterns of insular traveling waves varied reliably with behavioral states and also correlated with memory performance, we hypothesized that we would be able to use these signals to decode behavior at the single trial level. We trained multilayer neural networks, with cross-validation, for classifying pairwise behavioral states. We performed this decoding separately for each of the frequencies (**Methods, Figures 5, 6E**).

We first examined whether behavioral states can be decoded using the strength and propagation direction of the traveling waves. We found that behavioral states can be reliably decoded at the individual subject level and also at the group level across all frequencies (**Figures 5B-D**, *p’s* < 0.001, one-sided sign tests). Some behavioral states can be more reliably decoded than others (**Figures 5E-G**). For example, **Figure 5C** shows that for subject 7, the spatial wave patterns for the confidence and distractor periods were distinct from each other and hence were more easily distinguishable compared to those from the navigation and retrieval periods. Overall, distractor and navigation were more often distinguishable, which indicates that these behaviors generally exhibit similar traveling wave patterns (Dendrograms in **Figures 5E-G**).

Since the spatial patterns of insular traveling waves varied reliably with memory performance, we also hypothesized that we would be able to use these signals to decode each subject’s performance at the single trial level. Like before, we trained multilayer neural networks, with cross-validation, for classifying successful versus unsuccessful memory encoding, etc. This analysis revealed that memory performance can be reliably decoded across frequencies and subjects (**Figure 6E**, *p’s* < 0.001, one-sided sign tests).

### Spatially heterogeneous patterns of traveling waves in the anterior compared to the posterior insula

fMRI studies revealed that the anterior portion of the insula is functionally distinct from the posterior insula node (Menon, 2025). As a pivotal node of the salience network, along with the anterior cingulate cortex, the anterior insula is a functionally heterogeneous brain region which plays a critical role in regulating the engagement and disengagement of the default mode and frontoparietal control networks across diverse cognitive tasks (Cai et al., 2016; Menon, 2025; Sridharan et al., 2008). Furthermore, prior iEEG work on the salience network has revealed higher entropy, and therefore higher diversity, within anterior insula electrodes, compared to electrodes in the default mode and frontoparietal brain areas, highlighting both spatial and temporal heterogeneity within the anterior insula (Das & Menon, 2020, 2024).

Based on this, and similar to our analyses in the previous sections, we first examined the strength of traveling waves for the anterior insula compared to the posterior insula for each frequency and task condition. This analysis revealed that the anterior insula has reduced wave strength compared to the posterior insula across all task conditions and frequencies (**Figures 8A**, **S3**, *p’s* < 0.05, Wilcoxon sign-rank tests). Furthermore, this trend was observed in 9 out of the 10 subjects. Additionally, to rule out the possibility that the increased wave-strength of the posterior, compared to the anterior, insula could have been driven by a difference in signal power, we carried out control analyses comparing the power for the anterior and posterior insula. This analysis revealed that power does not differ between the anterior and posterior insula, in any of the frequency bands or in any of the task conditions (*p’s* > 0.05, Wilcoxon sign-rank tests).

**Figure 8:**
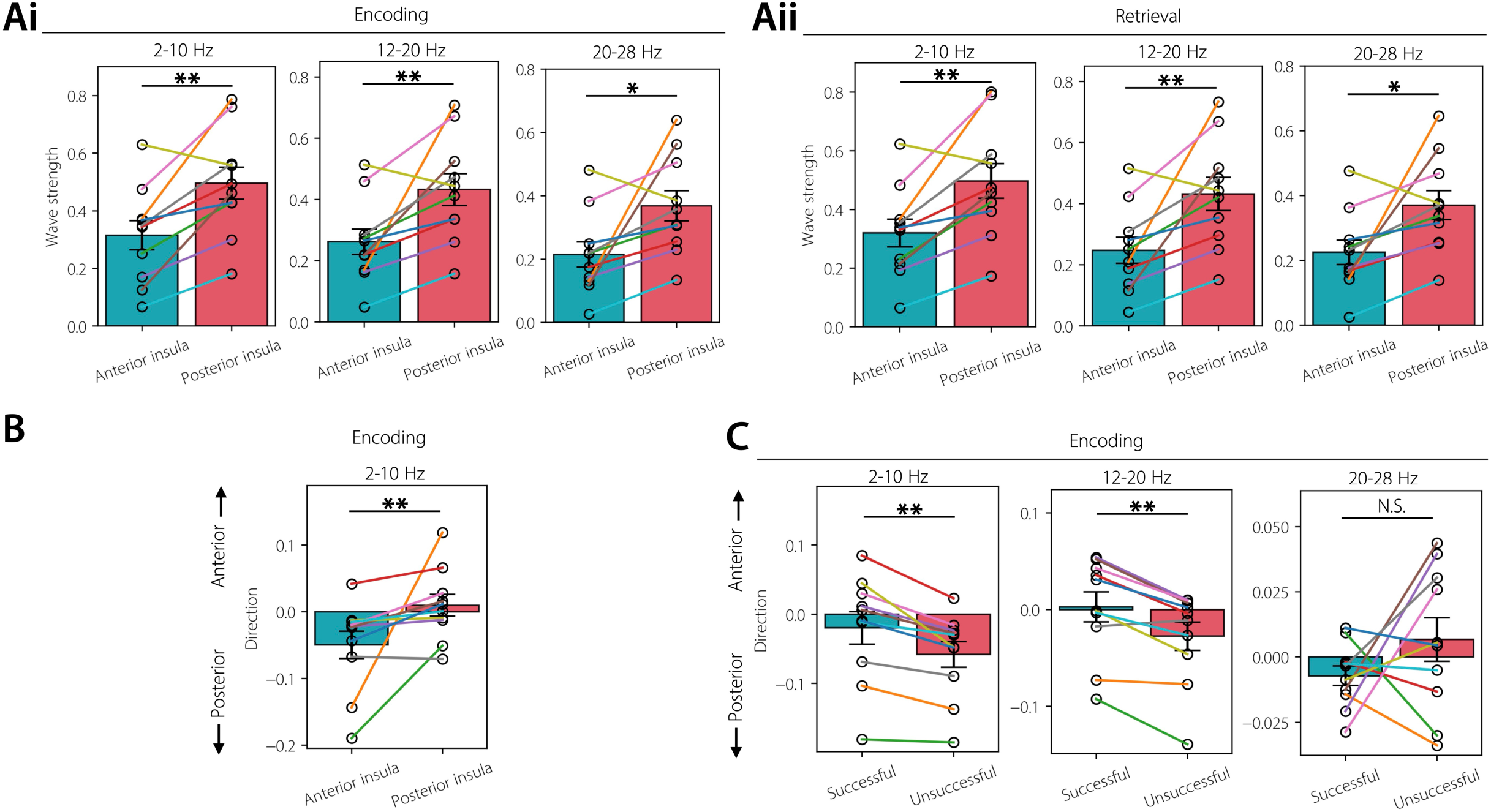
Wave strength and direction of insular traveling waves align with functional and anatomical parcellation of the insula. **(A)** The anterior insula has reduced wave strength compared to the posterior insula during the encoding **(i)** and retrieval periods **(ii)**, and across all frequencies. This finding was replicated across all other task conditions as well (**Figure S3**). This suggests higher spatial diversity and therefore, heterogenous information processing during memory formation within the anterior, compared to the posterior, insula. **(B) Waves in the anterior insula propagated posteriorly towards the posterior insula in the theta frequency during the encoding period.** Panel shows comparison of the anterior-posterior direction of the traveling waves in the anterior and posterior insula. This finding was also replicated across all other task conditions (**Figure S4**). **(C) Direction of the theta waves in the anterior insula is correlated with memory performance.** Comparison of the anterior-posterior direction of the traveling waves in the anterior insula for the successful compared to unsuccessful memory encoding, in each frequency. Posteriorly propagating traveling waves in the anterior insula get suppressed during successful compared to unsuccessful memory encoding in theta and low beta frequency bands. Error bars denote SEM across subjects. N=10. ** *p* < 0.01, * *p* < 0.05, N.S. Not significant (FDR-corrected).

Together, these findings suggest higher spatial diversity and heterogenous information processing during memory formation within the anterior insula.

We next examined the anterior-posterior direction of the traveling waves for the anterior compared to the posterior insula for each frequency and task condition. This analysis revealed that waves in the anterior insula propagated more posteriorly compared to the posterior insula in the theta frequency band, critically this finding was replicated across all task conditions (**Figures 8B**, **S4**, *p’s* < 0.05, Wilcoxon sign-rank tests). Propagation direction of the traveling waves did not differ between the anterior compared to the posterior insula in the low beta and high beta frequency bands (**Figure S4**, *p’s* > 0.05, Wilcoxon sign-rank tests). Together, these findings suggest that the insula is a control network in the low frequency theta band, directing information towards the posterior insula for putative detection and filtering of task relevant information across a diverse range of tasks (Cai et al., 2016; Menon, 2025; Sridharan et al., 2008). These findings also suggest that the anterior insula has directed influence not only on the default mode and frontoparietal networks (Das & Menon, 2024), but also on the posterior insula, a subregion within the insula.

To distinguish functional processes along the anterior-posterior axis, we further examined whether these posteriorly propagating waves in the anterior insula are also related to memory performance. This analysis revealed that the direction of these posteriorly propagating waves was suppressed during successful compared to unsuccessful memory encoding, both in the theta and low beta frequency bands (**Figure 8C**, *p’s* < 0.01, Wilcoxon sign-rank tests). Direction of these posteriorly propagating waves neither changed in any other task condition (*p’s* > 0.05, Wilcoxon sign-rank tests), nor in the posterior insula (*p’s* > 0.05, Wilcoxon sign-rank tests).

Suppression of these posteriorly propagating waves in the anterior insula during memory encoding suggests putative initiation of rapid switching dynamics by the anterior insula in response to shifting task demands for successful memory formation (Menon, 2025).

## Discussion

### Theoretical and functional implications of the insular traveling waves

Neuroimaging studies showed that the anterior insula, a core node of the salience network, is a functionally heterogeneous brain region that plays a foundational role in human cognitive processes. In particular, the insula interacts with other large-scale brain networks such as the medial default mode network and lateral frontal–parietal network, and in doing so regulates external stimulus-related cognition and monitoring internal mental processes (Menon, 2025). Interactions between these three networks, as described in the ‘triple-network’ model framework, therefore facilitate the performance of a diverse range of demanding cognitive tasks (Das & Menon, 2020, 2024; Sridharan et al., 2008).

Central to this model is the anterior insula. The anterior insula is consistently engaged during attentional tasks. Dynamic causal modeling of fMRI recordings suggests that human attentional processes are supported by the anterior insula exerting strong causal influences on the default mode and frontoparietal networks (Cai et al., 2016; Sridharan et al., 2008). The anterior insula is additionally considered critical because it is distinguished by the presence of von Economo neurons, which have a unique spindle shape and thick dendrites, and are theorized to be critical for effective cognitive control and interoception (Evrard, 2019; Nieuwenhuys, 2012). The anterior insula has a distinct set of projections to higher order cognitive regions, such as the orbitofrontal, entorhinal, amygdala, periamygdaloid, piriform, olfactory cortices, and the temporal pole, differentiating it from the posterior insula which is connected to primary and secondary sensory and motor cortices (Fudge et al., 2005; Höistad & Barbas, 2008; Mesulam & Mufson., 1985; Stefanacci & Amaral, 2002). By virtue of its higher causal influence over other brain areas, the anterior insula is well-positioned to dynamically engage and disengage with other brain areas, therefore facilitating rapid and efficient switching between the default mode and frontoparietal networks in response to shifting task demands (Menon, 2025).

In our study we found that the strength of traveling waves in the anterior insula is reduced compared to the posterior insula across diverse task conditions, therefore highlighting both spatial and temporal heterogeneity within the anterior insula. Higher spatial diversity in the anterior insula suggests diverse functional modules or sub-networks within the anterior insula each capable of a distinct, but specialized role for tackling a complex neural process such as memory formation and processing and therefore, may also enable the anterior insula for influencing bottom-up and top-down information processing loops. Spatial diversity reduces redundancy, increases flexibility, which consequently increases synergistic information flow, and multiple sub-regions within the anterior insula may jointly influence other brain areas. Spatial diversity ensures that if one signal path is impaired, others can compensate, effectively broadening the range of accessible states and making the insular cortex more flexible to shifting task demands. Thus, the distinctive traveling wave patterns in the anterior insula may be a vital part of this region’s distinctive functional role in cognition.

Furthermore, prior iEEG work on the salience network has revealed higher directed influence of the anterior insula on the default mode and frontoparietal networks as well as higher entropy, and therefore higher spatial and temporal diversity, within anterior insula electrodes compared to electrodes in the default mode and frontoparietal brain areas (Das & Menon, 2020, 2024).

Traveling waves with reduced strength in the anterior insula may therefore facilitate the anterior insula to dynamically engage and disengage with other higher order cognitive areas to initiate rapid switching dynamics for spatially heterogeneous and flexible information processing. In contrast, in the posterior insula the stronger traveling waves may be relevant for its less demanding and more spatially homogeneous information processing where it coordinates with simpler sensory brain areas.

In addition to distinguishing different functional states, we also found that insular traveling waves correlated with task performance, as the strength of traveling waves in the low beta band was greater for successful compared to unsuccessful memory encoding. This finding was reversed in the theta band. These performance-related traveling waves may be relevant for memory due to interactions between the insula and the prefrontal cortex, which plays a prominent role in controlling top-down information flow for transition and maintenance of latent neuronal ensembles into active representations (Das & Menon, 2021, 2022; Engel & Fries, 2010; Spitzer & Haegens, 2017). Top-down control requires coordination between multiple brain areas, and traveling waves provide a structured way to align the timing of excitability across diverse regions in space, allowing the control signal from insular cortex to propagate effectively. Since stronger waves mean more reliable phase gradients across space, this lets neurons in downstream areas know when to receive or ignore inputs — essentially enabling the insula to effectively route information according to the control goal. Strong insular traveling waves during successful memory encoding provides a robust carrier signal, potentially increasing the signal-to-noise ratio of the read-out of phase-modulated neurons, such as those involved in phase precession (O’Keefe & Recce, 1993). Thus, traveling waves may enable opening predictable windows of excitability, allowing higher-order control signals to selectively gate sensory processing in downstream brain areas. Therefore, increased wave strength during successful memory encoding in the beta band could be a putative mechanism that helps the insula to exert top-down influence for control of successful memory formation.

Conversely, low frequency theta waves are more related to hippocampal signaling of pattern completion associated with memory processing that is conveyed to multiple prefrontal regions (Eichenbaum, 2017), therefore decreased wave strength in the theta band during successful memory encoding might mark a network switch that disengages control networks in the insula and allows the default mode and hippocampal networks to drive memory formation. Suppression of insula traveling waves during successful encoding in the theta band could therefore reflect a functional disengagement of large-scale broadcasting in favor of more localized, content-specific processing. Because strong insular traveling waves might interfere with hippocampal–cortical binding of sparse patterns of neuronal spiking, suppression of waves could reduce this interference by allowing sensory and hippocampal circuits to dominate, which enables the hippocampus to support detailed, rich encoding and hippocampal–insular binding of specific details. Because, traveling waves also exist in the hippocampus during memory processing (Zhang & Jacobs, 2015) and also because brain stimulation studies have shown causal connectivity between the insula and the hippocampus (Huang et al., 2025), future studies may wish to identify links between subcortical waves with the insular waves that we focus on here.

The increase in wave strength during the unsuccessful memory trials compared to successful memory retrieval in the low beta frequency band is particularly intriguing. The insula is a critical brain region for detecting salient events, and therefore stronger traveling waves during unsuccessful memory retrieval suggests the role of insula in error processing, a salient event in itself, particularly in the context of conscious error perception (Ullsperger et al., 2010). Lesions to the anterior insular cortex are known to cause alterations in conscious error perception (Klein et al., 2013). Because the insula is known for controlled retrieval (Menon, 2011), increased traveling waves can broadcast an error signal across brain-wide networks, therefore initiating cognitive control adjustments and a shift in attentional resources to focus on the erroneous events and suppressing irrelevant activity, and prevent future similar mistakes during retrieval to improve performance. Increased insular waves during unsuccessful memory retrieval recapitulates insula’s role as a control hub, initiating propagation of corrective information across large-scale memory networks. Strong insular wave’s phase gradients provide a temporal scaffold for aligning distributed regions, while its propagation enables the wide dissemination of error and control signals needed for reorienting memory search.

### Possible mechanisms for the complex spatial patterns of insular traveling waves

In the traditional neural hierarchy, sensory-related neural activity usually travels in a feedforward way whereas neural activity related to higher-order cognitive processes such as memory retrieval feeds “backward” from frontal cortices to reinstate neural activity in other brain regions (Rabinovich et al., 2012). Our earlier study (Mohan et al., 2024) had shown a role for these forward and backward planar traveling waves. However, recently we have shown a role for complex spatial patterns of traveling waves on the surface of the cortex, such as spirals and source/sink waves, during memory (Das et al., 2025). Our findings of complex spatial patterns of insular traveling waves suggest an additional new type of spatial organization of electrophysiological activity in the cortex, complementing the feedforward-feedback cortical hierarchy, converging with related findings of complex wave patterns in rodents and non-human primates (Bhattacharya et al., 2022; Liang et al., 2023) as well as in humans (Muller et al., 2016), playing a critical role in modulating behavior. Since traveling waves are known to be closely associated with spiking activity of neurons (Davis et al., 2020), their propagation putatively reflects packets of discrete neuronal activity going along with the wave peak/trough and sequentially scanning distributed insular areas to transiently reorganize functional connectivity between them, to represent complex behaviors of human memory representations (Eichenbaum, 2000; Mesulam, 1990). Because spiral waves revisit the same brain areas in multiple cycles, it suggests that these waves may be relevant for dynamically strengthening functional connectivity between large-scale neuronal assemblies for efficient memory processing, similar to the rotational waves observed during sleep spindles (Muller et al., 2016). Similarly, the insular source waves that we observed indicate the presence of a small local group of neuronal assemblies that dominate information flow by routing their information outward in a direction where they flow towards widespread brain areas. The other complex, heterogeneous spatial patterns of insular traveling waves that we observed putatively route information in more flexible ways by propagating in several directions to rapidly reorganize functional connectivity between neuronal assemblies, to distinguish behavioral states. Our results on insular waves suggest that this additional new hierarchy also holds true for deep neocortical structures such as the insula, and not just for neural activity on the surface of the cortex (Das et al., 2025).

The direction of wave propagation in the insula may not only reflect the spatial routing of information, but also actively shape it, through neural coding and synaptic plasticity.

Specifically, the direction of wave propagation — such as anterior-to-posterior movement across the insula — may temporally structure neural activity in a way that promotes the formation of new neural assemblies along the wave’s path (Buzsáki, 2010). This organization could be mediated by phase coding, whereby the timing of neuronal firing relative to the wave cycle supports precise spike timing and synchrony (Bi & Poo, 1998). Based on Hebbian principles, therefore, over time, such temporally coordinated activation may induce long-term potentiation, reinforcing synaptic connections in the direction of the wave and sculpting functional networks within the insula (Song et al., 2000). Furthermore, these waves may entrain downstream regions, causing corresponding assemblies to form or reorganize in accordance with the temporal structure of insular outputs. This mechanism offers a novel account of how dynamic, wave-based neural activity in the insula may drive both local and network-level plasticity.

Going forward, it will be important to understand how complex patterns of insula traveling waves emerge and what they mean for underlying neuronal computations. One theoretical framework that could explain how brain oscillations show traveling waves with complex spatial patterns is neural models based on weakly coupled oscillators (Bhattacharya et al., 2021; Sato, 2022). These models hypothesize that complex patterns of waves can be generated locally based on the initial spatial activation of neurons, where each neuron is connected to a few of its neighbors, with distance dependent axonal delays in the order of conduction along unmyelinated horizontal fibers (Davis et al., 2021; Destexhe, 1994; Ermentrout & Kleinfeld, 2001). These locally generated waves can propagate across widespread regions between locally connected neurons and interact with other locally generated waves, to generate complex patterns of propagating oscillations (Huang et al., 2010; Schiff et al., 2007). A key determinant of the shape of these wave patterns is the presence of local shifts in the amplitude and frequency of local oscillations, which posits that waves tend to propagate away from the cortical locations with the fastest intrinsic oscillation frequencies, following a gradient to the locations with the slowest oscillations (Kopell & Ermentrout, 1990). Therefore, even though the insular waves remain spatiotemporally stable within an individual participant, variations in the local neuronal connections along with the heterogeneity in the amplitude and frequency of local oscillations and their spatial locations across participants can result in a wide range of complex wave patterns across the insula of individual participants (Kopell & Ermentrout, 1990).

## Conclusions

The diverse shapes of insular traveling waves that we observed during the various task periods help the insula flexibly initiate control mechanisms, regulating the transition of information within the insula across different memory stages and also towards other memory-related brain areas such as the hippocampus as well as sensory-related brain areas. Specifically, the traveling waves in the anterior insula travelled posteriorly towards the posterior insula in theta frequency, suggesting that the anterior insula is a control network, directing information towards the posterior insula for putative detection and filtering of task relevant information. Crucially, these posteriorly moving theta waves in the anterior insula were suppressed during successful memory formation compared to when memory formation was unsuccessful, suggesting putative initiation of rapid switching dynamics by the anterior insula in response to shifting task demands for successful memory formation (Menon, 2025). Therefore, our findings of insular traveling waves provide a novel neurophysiological model for understanding the anterior insula as a dynamic switch, to flexibly encode sensory information into short-term memory for efficient, goal-oriented information routing during complex behaviors in human memory processing. Because a lack of efficient switching dynamics of the insula with other cortical areas appears in neurological disorders such as depression (Teckentrup et al., 2025), insular traveling waves could be a biomarker for investigating dysfunctions in neurological disorders.

## Materials and Methods

### Human subjects

We examined direct brain recordings from 10 patients with pharmaco-resistant epilepsy who underwent surgery for removal of their seizure onset zones. Only patients which had a minimum of 6 insular contacts were included in our analysis since estimating traveling waves with very low number of electrodes renders unreliable results (Mohan et al., 2024). All patients consented to having their brain recordings used for research purposes and all research was approved by Institutional Review Boards. The patients (N=10, 6 females, minimum age = 19, maximum age = 59, mean age = 40.6, see below for details) were part of a larger data collection initiative that can be downloaded from the UPENN-RAM consortium (URL: http://memory.psych.upenn.edu/RAM), shared by Kahana and colleagues (Jacobs et al., 2016).

These data were recorded at eight hospitals: Thomas Jefferson University Hospital; University of Texas Southwestern Medical Center; Emory University Hospital; Dartmouth College Hospital; University of Pennsylvania Hospital; Mayo Clinic; National Institutes of Health; and Columbia University Hospital. Prior to data collection, research protocols and ethical guidelines were approved by the Institutional Review Board at the participating hospitals and informed consent was obtained from the participants and guardians (Jacobs et al., 2016).

### Electrophysiological recordings and preprocessing

Patients were implanted with different configuration of electrodes based on their clinical needs, which included both electrocorticographic surface grid and strips as well as depth electrodes. In this work, we only examined the depth electrodes (contacts spaced 5–10 mm apart) implanted inside the insular cortex. Anatomical localization of electrode placement was accomplished by co-registering the postoperative computed CTs with the postoperative MRIs using FSL (FMRIB (Functional MRI of the Brain) Software Library), BET (Brain Extraction Tool), and FLIRT (FMRIB Linear Image Registration Tool) software packages. Preoperative MRIs were used when postoperative MRIs were not available. From these images, we identified the location of each recording contact on the CT images and computed the electrode location in standardized Talairach coordinates. We used the insula atlas in the Brainnetome atlas (Fan et al., 2016) to demarcate the insula. This atlas also consists of the three short dorsal gyri in the anterior insula and the two long gyri in the posterior insula. Total number of electrodes implanted in the insular cortex was 150 across all patients, with 83 electrodes in the anterior and 67 electrodes in the posterior insula. The number of electrodes in the anterior insula did not differ from those in the posterior insula (*p* > 0.05, binomial test), suggesting that the number of electrodes has limited impact on the anterior versus posterior results reported here.

Original sampling rates of these intracranial EEG (iEEG) signals were 500 Hz, 1000 Hz, and 1600 Hz. These iEEG signals were downsampled to 500 Hz, if the original sampling rate was higher, for all subsequent analysis. We used common average referencing (iEEG electrodes re-referenced to the average signal of all electrodes in the same subject), similar to our previous studies on traveling waves (Das et al., 2022; Das et al., 2025). Line noise (60 Hz) and its harmonics were removed from the iEEG signals. iEEG signals were filtered in the theta (2-10 Hz), low beta (12-20 Hz), and high beta (20-28) frequency bands. We filtered our iEEG signals in these relatively broad frequency bands rather than a narrowly defined frequency band because recent electrophysiology studies in nonhuman primates have suggested that broadband field potentials activity, rather than narrowband, governs wave dynamics in the brain (Davis et al., 2021; Davis et al., 2020). For filtering, we used a fourth order two-way zero phase lag Butterworth filter throughout the analysis.

### Treasure Hunt spatial episodic memory task

The patients performed multiple trials of a spatial memory task in a virtual reality environment (Miller et al., 2018) on a laptop computer at the bedside. They controlled their translational and rotational movements through the virtual environment with a handheld joystick. In each task trial, subjects explored a rectangular arena on a virtual 3D beach to reach treasure chests that revealed hidden objects, with the goal of encoding the location of each item encountered (**Figure 1A**). The virtual beach (100 × 70 virtual units, 1.42: 1 aspect ratio) was bounded by a wooden fence on each side. The locations of the objects changed over the trials, but the shape, size and appearance of the environment remained constant throughout the sessions. The task environment was constructed so that the subject would perceive one virtual unit as corresponding to approximately 1 foot in the real world. Subjects viewed the environment from the perspective of cycling through the environment and the elevation of their perspective was 5.6 virtual units. Each end of the environment had unique visual cues to help the subjects navigate.

Each trial of the task begins with the subject being placed on the ground at a randomly selected end of the environment. The subject initiates the trial with a button press, then navigates to a chest using a joystick. Upon arrival at the chest, the chest opens and either reveals an object, which the subject should try to remember, or is empty. The subject remains facing the open chest for 1.5 sec (*encoding period*) and then the object and chest disappear, which indicates that the subject should navigate (*navigation period*) to the next chest that has now appeared in the arena. Each trial consists of four chests; two or three (randomly selected, so that subjects could not predict whether the current target chest contained an object, which served to remove effects of expectation and to encourage subjects to always attend to their current location as they approached a chest) of the chests contain an object, and the others are empty. Each session consists of 40 trials, and in each session, subjects visit a total of 100 full chests and 60 empty chests. The chests are located pseudorandomly throughout the interior of the environment. After reaching all four chests of a trial, subjects are transported automatically so that they have an elevated 3/4 overhead perspective view of the environment at a randomly selected end of the environment. They then perform a distractor task (*distractor period*), a computerized version of the “shell game”, before entering the *retrieval period*. During recall, subjects are cued with each of the objects from the trial in a random sequence and asked to recall the location of the object.

In each recall period, they first indicate their confidence (*confidence period*) to remember the location of the object (“high”, “medium”, or “low”). Next, they indicate the precise location of the object by placing a cross-hair at the location in the environment that corresponds to the location of the cued item. After the location of each object of the trial is indicated, the feedback stage (*feedback period*) of each trial begins. Subjects are shown their response for each object and given feedback on their response accuracy, following a point system where they receive greater rewards for accurate responses. A response is considered correct if it is within 13 virtual units of the true object location. Mean accuracy across subjects was ∼33%.

We analyzed the 1.5-sec long trials from the encoding periods of the task. For the navigation periods, we analyzed 1.5 sec long segments approximately corresponding to the middle of the navigation trial. Similarly, for the distractor periods, we analyzed 1.5 sec long time segments approximately corresponding to the middle of the distractor trial. For the confidence periods, we analyzed 1.5 sec recording immediately following the presentation of the visual cues. For the retrieval periods, we analyzed 1.5 sec immediately prior to the retrieval of the objects. For the feedback periods, we analyzed 1.5 sec time segments immediately following the feedback.

The 10 subjects completed 17 sessions of the memory task in total. On an average, each subject completed ∼ 145 encoding trials, ∼ 90 confidence trials, ∼ 62 navigation trials, ∼ 78 retrieval trials, ∼ 62 distractor trials, and ∼ 66 feedback trials, in each session of the spatial task. Such a relatively large number of trials for the different task periods ensured a high degree of statistical power to yield robust results.

### Identification of traveling waves

We first filtered the iEEG signals in the theta, low beta, and high beta frequency bands. We identified and characterized traveling waves in each of these frequency bands separately.

Quantitatively, a traveling (phase) wave can be measured as a set of simultaneously recorded neural activity whose instantaneous phases vary systematically with the locations of the recording electrodes. To identify these traveling waves we used a localized circular-linear regression approach, assuming that the relative phases of the iEEG signals exhibit a linear relationship with electrode locations *locally* (Das et al., 2022; Das et al., 2025). This locally circular-linear fitting of phase-location can detect complex patterns (Ermentrout & Kleinfeld, 2001; Muller et al., 2016) of traveling waves in addition to planar traveling waves.

To identify traveling waves from the phases, we first measured the instantaneous phases of the signals from each electrode of a given cluster by applying a 4^th^-order Butterworth filter at each frequency band separately (bandwidth [f_p_ ×.85, f_p_ / .85] where f_p_ is the peak frequency). We used the Hilbert transform on each electrode’s filtered signal to extract the instantaneous phase.

Next, we used circular statistics to measure the spatial propagation of phase between neighboring electrodes to identify traveling waves at each time point (Fisher, 1993). To simplify visualizing and interpreting the data, we first projected the 3-D Talairach coordinates of insular electrodes to a 2-D plane using principal component analysis (PCA).

To identify traveling waves, we used a series of two-dimensional (2-D) localized circular–linear regression that were fit to model the direction of wave propagation in a local *subcluster* in the 25-mm neighborhood of each electrode. The local regression determines the direction of local wave propagation in the subcluster surrounding each electrode, by measuring whether the local phase pattern varies linearly with the electrodes’ coordinates in 2-D. Thus, this regression fits a single direction of wave propagation locally around each electrode at each moment, and these measures are then subsequently combined across subclusters to identify larger-scale spatial patterns of propagation.

In this regression, for each nearby electrode, we first identified the neighboring electrodes that were located nearby, constituting a sub-cluster. Here, let *x_i_* and *y_i_* represent the 2-D coordinates and *θ_i_* the instantaneous phase of the *i*th electrode in a sub-cluster. We used a 2-D circular-linear model

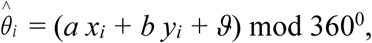

where 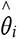 is the predicted phase, *a* and *b* are the phase slopes corresponding to the rate of phase change (or spatial frequencies) in each dimension, and *ϑ* is the phase offset. We converted this model to polar coordinates to simplify fitting. We define *α* = atan2(*b, a*) which denotes the angle of wave propagation and 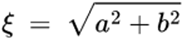 which denotes the spatial frequency. Circular–linear models do not have an analytical solution and must be fitted iteratively (Fisher, 1993). We fitted *α* and *ξ* to the distribution of phases at each time point by conducting a grid search over *α* ∈ [0°, 360°] and *ξ* ∈ [0, 18]. Note that *ξ* = 18 corresponds to the spatial Nyquist frequency of 18^0^/mm corresponding to the spacing between neighboring electrodes of 10 mm.

In order to keep the computational complexity tractable, we used a multi-resolution grid search. We first carried out a grid search in increments of 5^0^ and 1^0^/mm for *α* and *ξ*, respectively. The model parameters (*a = ξ*cos(*α*) and *b= ξ*sin(*α*)) for each time point are fitted to most closely match the phase observed at each electrode in the sub-cluster. After having relatively coarse estimates of *α* and *ξ*, we then carried out another grid search in increments of 0.05^0^ and 0.05^0^/mm around a ± 2.5^0^ and ± 0.5^0^/mm neighborhood of the coarse estimates of *α* and *ξ*, respectively, to have refined estimates of *α* and *ξ*. We computed the goodness of fit as the mean vector length 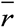 of the residuals between the predicted (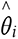) and actual (*θ_i_*) phases (Fisher, 1993),

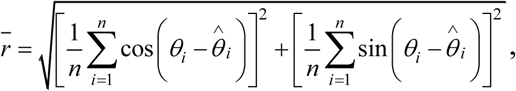

where *n* is the number of electrodes in the sub-cluster. The selected values of *α* and *ξ* are chosen to maximize 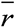. This procedure is repeated for each sub-cluster. To measure the statistical reliability of each fitted traveling wave, we examined the phase variance that was explained by the best fitting model. To do this, we computed the circular correlation *ρ_cc_* between the predicted (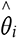) and actual (*θ_i_*) phases at each electrode:

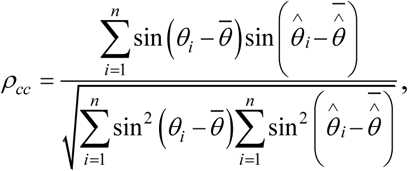

where bar denotes averaging across electrodes. We refer to *ρ*^2^*_cc_* as the wave strength (Das et al., 2022) as it quantifies the strength of the traveling wave (note that *ρ*^2^*_cc_* has been referred to as the phase gradient directionality (PGD) in some prior studies (Muller et al., 2016; Zhang et al., 2018)). Other features of traveling waves such as the wavelength (2π/spatial frequency) and the speed (wavelength × frequency) can be readily derived from the parameters of the above 2-D model. Note that traveling waves in some prior studies were detected and analyzed by calculating the spatial gradient of the phases of the iEEG recordings (Halgren et al., 2019; Muller et al., 2016), however, phase gradients can only be calculated in two directions (forward and backward), so only a subset of neighboring electrodes of a given electrode are included in these analyses of spatial gradient. Since our approach directly includes all possible neighboring electrodes (termed as a sub-cluster in our analysis) in the circular-linear regression model, thus calculating phase gradients in all possible directions, it results in a more efficient estimate of the traveling waves parameters.

### Identification of stable epochs

Since a traveling wave is composed of phase patterns that vary relatively smoothly across space and time, we sought to characterize the spatiotemporal stability of the traveling waves that we detected in our localized circular-linear regression approach above. We defined *stability* as the negative of the mean (across all insular electrodes in a given subject) of the absolute values of the difference between the direction and strength of traveling waves observed at consecutive time-points, defined as

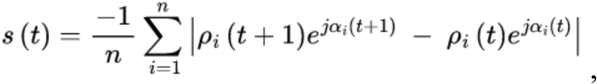

where, *ρ_i_* denotes the strength of the traveling wave for the *i*^th^ electrode, *α_i_* denotes the direction of the traveling wave for the *i*^th^ electrode, *n* denotes the number of insular electrodes, *s* denotes stability, *t* denotes time, and *j* denotes square root of minus one. We repeated this procedure for each pair of consecutive time-points and for each trial of the encoding, retrieval, navigation, etc. periods. Stability for each trial was z-scored. We identified stable epochs as those for which all stability values were above the mean of the stability values for a given trial. We ran our stability analysis across all trials (**Figures 2B, C**).

Previous research in rabbit field potential recordings (Freeman & Rogers, 2002; Freeman & Schneider, 1982) have found spatial patterns of theta traveling waves last ∼80-100 msec in duration. Moreover, large-scale, whole-brain computational modeling in humans using neural field theory have shown that spatiotemporally stable traveling waves last ∼50-60 msec in duration (Roberts et al., 2019). Therefore, we focused our analyses on the stable epochs that were more than 50 msec long for further analysis; as there was evidence that shorter wave patterns may be less stable (Roberts et al., 2019). Overall, ∼ 46% of stable-epochs were long enough to be included based on this criteria, yielding ∼ 5 stable-epochs per trial.

### Identification of modes using complex independent component analysis (CICA)

We next used a large-scale dimensionality reduction framework to measure the spatial patterns of wave propagation across all insular electrodes for each subject. This framework identified the different types of spatial wave patterns that formed traveling waves in each individual subject and modeled how they summed to determine the signal on each epoch. We thus applied complex ICA (CICA) to the data on each epoch. Within each stable epoch identified as described above, the direction and strength of the traveling waves remain almost consistent across time-points.

Thus, we averaged the direction and strength of the traveling waves within each stable epoch to find one spatial wave pattern associated with each stable epoch, which was input to the CICA algorithm.

Next, to analyze how the spatial patterns of waves changed over time and related to behavior, we concatenated the wave patterns (i.e., direction and strength) for all stable epochs across encoding, retrieval, navigation, etc. periods into a single matrix and then passed this matrix as input to the complex version of the independent component analysis (CICA) (Fu et al., 2015; Li & Adalı, 2010). We used this complex version of the ICA, as compared to real ICA, to incorporate the 2-D directions of the traveling waves, weighted by the strength (*ρ* cos (*α*) and *ρ* sin (*α*)), defined for each insular electrode. We then extracted the independent activation functions (or, weights) (each activation function corresponds to one of the stable epochs) and the corresponding modes (**Figures 2E, F**) as the output from the CICA (Fu et al., 2015; Li & Adalı, 2010). Multiplication of each of these modes with the mean of the weights across all stable epochs corresponding to that specific mode results in a unique wave pattern associated with that mode. At the individual epoch level, a higher CICA weight for that epoch corresponding to a specific mode indicates higher representation of that wave pattern in that specific epoch and a lower CICA weight for an epoch corresponding to a specific mode indicates lower representation of that wave pattern in that specific epoch (Das et al., 2025). Moreover, the higher the variance explained by a given mode, the higher will be its representation across the trials. Generally, we observed more complex wave patterns in higher order modes (modes 2 and beyond), which suggests that complex wave patterns explain relatively less variance in the data compared to the planar waves. In this way, we can extract the CICA weights for each of the encoding, retrieval, navigation, etc. periods and directly compare them statistically; see ***Statistical analysis*** section for details. In our previous work, we have extensively validated the application of the CICA procedure to analyzing traveling waves to confirm its sensitivity to extract expected wave patterns and demonstrate specificity to reject negative results in the absence of such patterns (Das et al., 2025).

### Statistical analysis of the relation between insular traveling waves and behavior

i. *Robustness of traveling waves at the individual epoch level*

We conducted surrogate analysis to test the significance of the estimated stable epochs (see ***Identification of stable epochs*** section above) and whether the observed stable epochs are beyond chance levels. For each time-point, we shuffled the trial labels so that the temporal contiguity within a given trial is destroyed and also the electrodes, so that the spatial pattern for the corresponding behavioral state is destroyed, and then ran the stable epoch analysis using identical methodology as above. In this way, we built a surrogate distribution by aggregating all time-points corresponding to these shuffled stable epochs against which we then compared the aggregated time-points from the empirical stable epochs (*p* < 0.05).

(ii) *Identifying significant modes in CICA*

To estimate the number of significant modes in the CICA output for each cluster, we compared the variance explained for each mode with the theoretical variance threshold 100/*n*, where *n* is the number of insular electrodes in an individual subject. This theoretical variance corresponds to the variance of each mode if the total variance (100%) is equally distributed among all modes.

Because of the spatial structure of traveling waves, some modes will explain more variance compared to the other modes in the empirical data. We additionally shuffled the locations of the electrodes in each subject and recalculated the variance distribution across modes and confirmed that the shuffled variance for all the modes converged to the theoretical variance threshold of 100/*n*.

(iii) *Multivariate analysis of variance (MANOVA)*

To identify how traveling wave patterns related to behavior, we directly compared the weights estimated from the CICA procedure above between the encoding, retrieval, navigation, etc. periods, for each mode using multivariate analysis of variance (MANOVA). MANOVA was used because the weights were complex. We statistically distinguished weights corresponding to different behavioral states using the following model: *Real + Imag ∼ States*, where *Real* and *Imag* are the real and imaginary parts of the weights respectively and *States* are encoding, retrieval, navigation, etc. We used this model for each mode and applied FDR-corrections for multiple comparisons (*p* < 0.05) across all modes, frequencies, and subjects. Statistical significance from the MANOVA would indicate that traveling waves shift their direction and/or strength to form distinct directional patterns which can distinguish different behavioral states.

We designated a subject to be significant if at least one of the modes from the CICA in at least one of the frequency bands (theta/low beta/high beta) showed statistical significance in MANOVA. Moreover, we also repeated our MANOVA analysis by shuffling the labels of the different behavioral states such as encoding, retrieval, etc. and built a histogram from these shuffles (f-statistics) against which we compared the empirical data. This analysis revealed that all our MANOVA results were still statistically significant (*p’s* < 0.05).

(iv) *Wilcoxon sign-rank tests*

For comparing wave strength, direction, and power for memory performance (successful vs. unsuccessful) and feedback (good vs. bad) across subjects, we used Wilcoxon sign-rank tests (*p* < 0.05). Similarly, for comparing wave strength, direction, and power for anterior versus posterior insula, we also used Wilcoxon sign-rank tests (*p* < 0.05). All results were subsequently FDR-corrected for multiple comparisons across frequencies and task conditions. For comparing the wave strength and direction for task conditions, we used the Kruskal-Wallis test (*p* < 0.05). These results were then subsequently FDR-corrected for multiple comparisons across frequencies.

### Decoding analysis of the relation between insular traveling waves and behavior

Our final goal was to test whether we could robustly decode the behavioral states from the insular traveling wave patterns. Decoding behavioral states in our datasets is a multiclass classification problem. We converted this problem into a series of binary classification tasks, as these can be solved straightforwardly with various multivariate algorithms. We trained multilayer neural networks (Bernardi et al., 2020), with cross-validation, for classifying pairwise behavioral states (for example, encoding versus retrieval), training a neural network classifier for each frequency and subject separately. We used a PyTorch-based five-layer neural network for this binary classification problem (Paszke et al., 2019). We used the extracted weights from all the modes from the CICA procedure as features for training our neural network classifiers.

The multilayer network architecture comprised an input layer, three hidden layers (32, 16, and 8 neurons respectively), and an output layer with a sigmoid activation function, similar to the neural network architectures previously used for classifying behavioral states (Bernardi et al., 2020). The inputs for all our decoding analysis were the activation functions or the weights since for a given mode, these were changing across different task periods. Please note that for a subject with “N” number of insular electrodes and hence “N” number of dimensions in CICA, a 2N dimensional neural network was trained (the factor 2 comes from the fact that the weights are complex numbers and hence real and imaginary parts were considered separately). ReLU activation functions were applied to the hidden layers to introduce nonlinearity and improve generalization (Dahl et al., 2013). To further enhance generalization, dropout regularization was implemented after each hidden layer (Srivastava et al., 2014). For model optimization, we used Binary Cross Entropy as the loss function, ideal for binary classification tasks (Ruby & Yendapalli, 2020), and the Adam optimizer during training (Paszke et al., 2019).

To ensure robust optimization and generalization, we employed cross-validation for iterative training over multiple epochs. For each pair of behavioral states, we first rebalanced the data such that we have an equal number of epochs per behavioral state (van Gerven et al., 2013). We used a five-fold cross-validation technique in which the network model was fitted on 80% of the data and then its performance was tested on the remaining 20%. We used a subsampling procedure where each epoch was randomly assigned to a particular fold, subject to the constraint that all behavioral states are evenly represented (van Gerven et al., 2013), and then averaged the network decoding accuracies across folds.

To access group level statistical significance for the decoding accuracy results, we used one-sided sign tests versus chance (*p* < 0.05). Moreover, we also carried out additional statistical analysis by shuffling the labels of the behavioral states such as encoding, retrieval, etc. and then running the same decoding analysis on this shuffled data to generate a distribution of shuffled data against which the actual decoding performance can then be compared. This analysis revealed that our decoding results are statistically significant across all subjects (all *p’s* < 0.05), suggesting that structure of wave activity is the source of the classification accuracy.

Please note that all our traveling waves analysis were carried out for each subject separately since each subject showed specific patterns of wave strength and directions, which differed across subjects. Due to the variability of the strength and directions of the traveling waves, our decoder was also run in each subject separately rather than collapsing across subjects.

## Acknowledgements

This research was supported by an NSF CRCNS grant to J.J.

## Code availability

The codes for traveling waves analysis are available at https://github.com/anupdas777/complex_traveling_waves/tree/main.

---

**Supplementary Figure 1:**
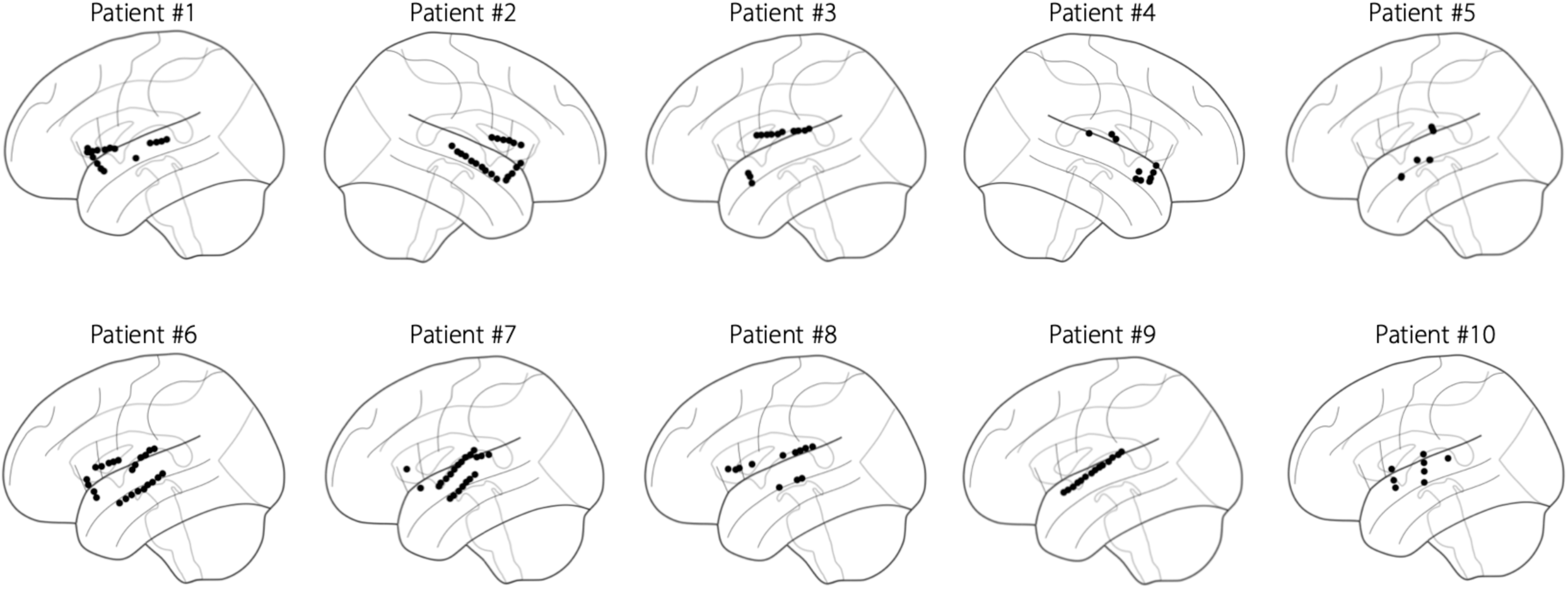
Insular electrode locations in the 10 subjects.

**Supplementary Figure 2:**
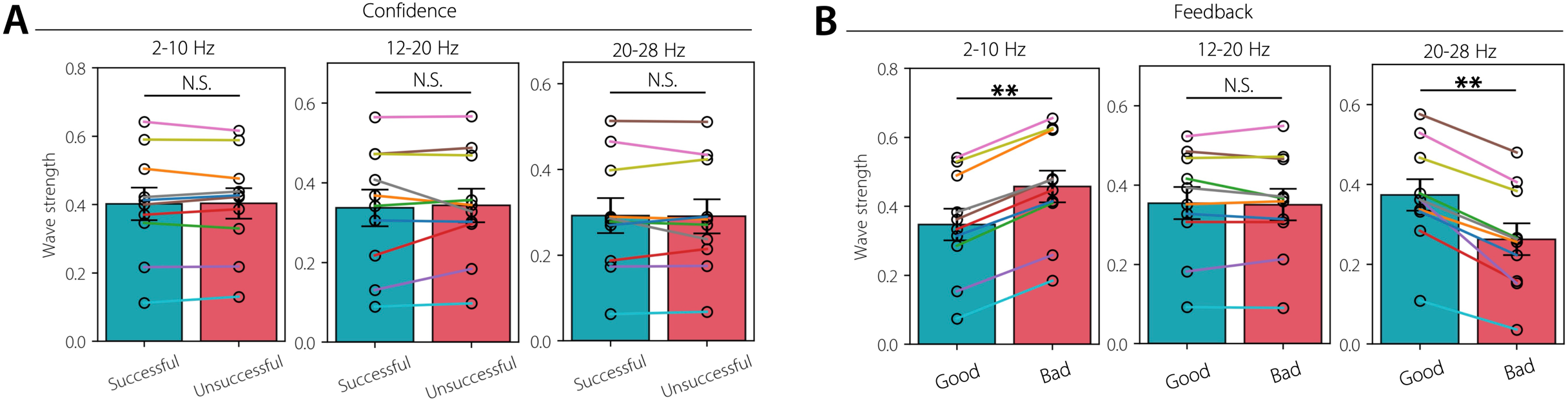
**(A) Wave strength of insular traveling waves for successful vs. unsuccessful memory confidence in each frequency band.** Error bars denote SEM across subjects. N=10. **(B) Same as in A, but for the feedback period.** Wave strength increased for good compared to bad feedback in the 20-28 Hz frequency. This pattern was reversed in the 2-10 Hz frequency. ** *p* < 0.01, N.S. Not significant (FDR-corrected).

**Supplementary Figure 3:**
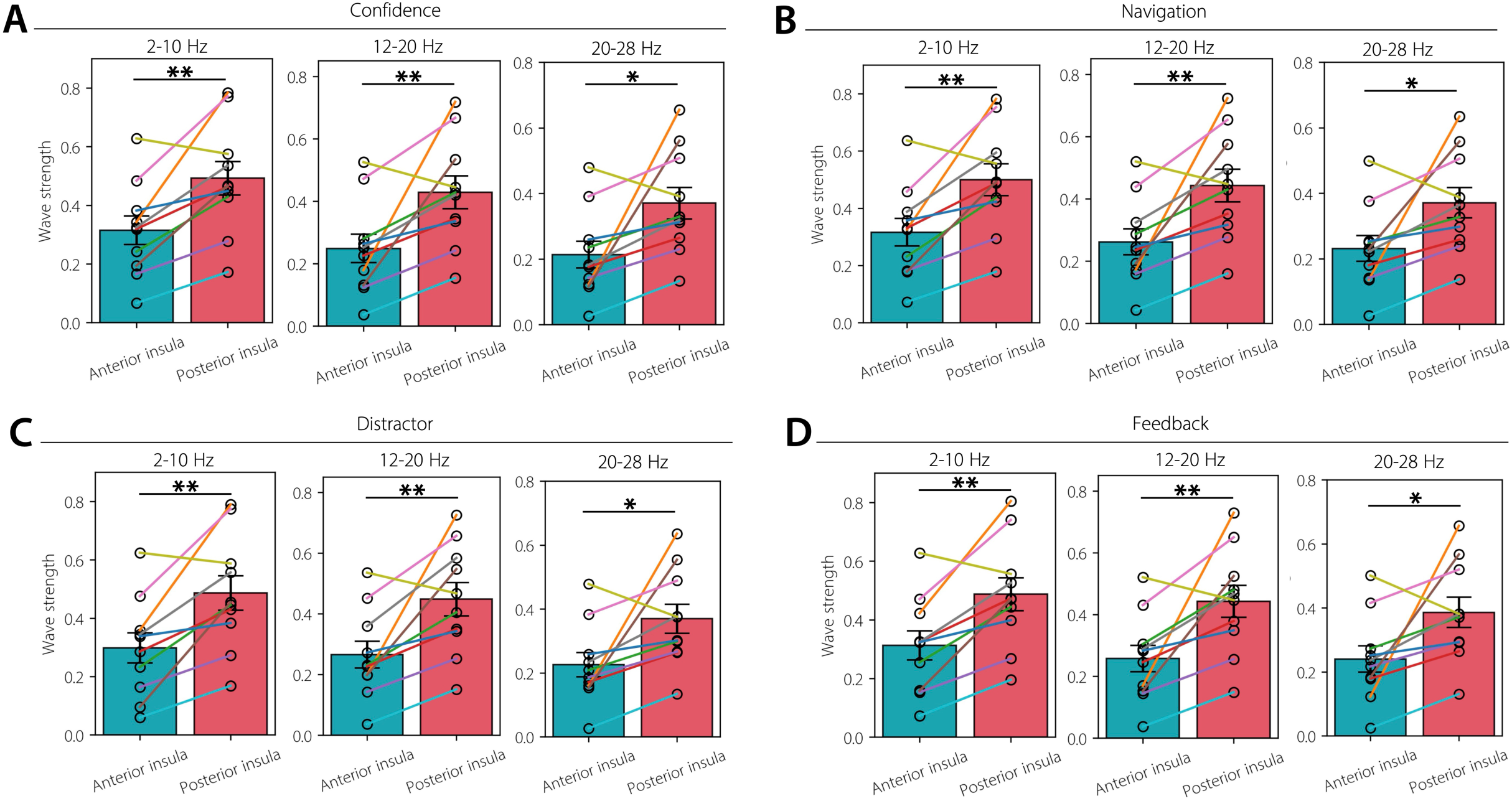
**Wave strength of insular traveling waves aligns with functional and anatomical parcellation of the insula. (A-D)** The anterior insula has reduced wave strength compared to the posterior insula across all task conditions and frequencies. Error bars denote SEM across subjects. N=10. ** *p* < 0.01, * *p* < 0.05 (FDR-corrected).

**Supplementary Figure 4:**
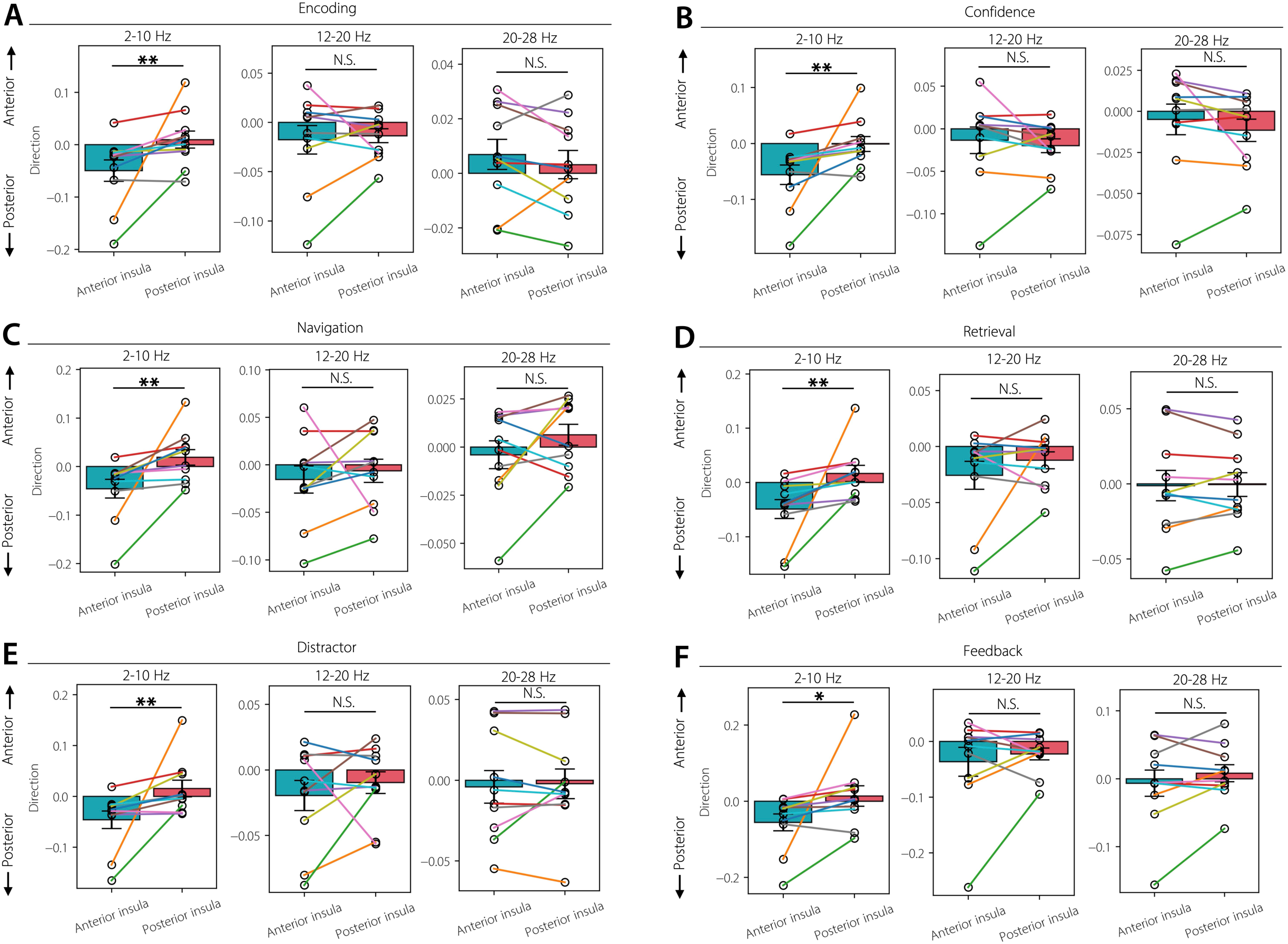
**Waves in the anterior insula propagated posteriorly towards the posterior insula in the theta frequency. (A-F)** Comparison of the anterior-posterior direction of the traveling waves in the anterior and posterior insula in each task condition and frequency. Error bars denote SEM across subjects. N=10. ** *p* < 0.01, * *p* < 0.05, N.S. Not significant (FDR-corrected).

